# Loss of MYSM1 deubiquitinase catalytic activity protects against MYC-driven B cell lymphoma via tumor intrinsic effects and indirect modulation of antitumor immunity

**DOI:** 10.64898/2026.01.19.700060

**Authors:** Dania Shaban, Viktoria Plackoska, Yue Liang, Nay Najm, Francis Robert, Sidong Huang, Anastasia Nijnik

## Abstract

**Background:** MYC is an oncogenic transcription factor that is over-expressed, amplified, or otherwise dysregulated in over 50% of all cancers. This includes over 10% of diffuse large B-cell lymphomas (DLBCL), where *MYC* translocations are associated with a poor therapy response and inferior prognosis for the patients. However, because MYC lacks ligand-binding or catalytic domains, it is a highly challenging drug target, and there is a wide interest in novel approaches to inhibit MYC oncogenic functions. MYSM1 is a chromatin-binding deubiquitinase (DUB) that promotes gene expression by catalytically removing the histone H2AK119ub epigenetic mark. In recent work, we demonstrated that MYSM1 acts in cooperation with MYC to sustain the expression of oncogenic transcriptional programs in hematopoietic cells, identifying MYSM1 as a potential therapeutic target for MYC-driven malignancies.

**Results:** Here, we show for the first time that the loss of MYSM1 DUB catalytic activity, without the loss of MYSM1 protein expression, is sufficient to protect against MYC-driven lymphoma in murine models. We characterize the impact of MYSM1 loss-of-function on tumor cell physiology and on antitumor immunity, examining the tumor-intrinsic and the immune cell-mediated mechanisms involved in the protection against the disease. Leveraging human cancer genome databases, we provide first evidence linking MYSM1 loss-of-function to reduced fitness of human lymphoma cell lines in culture and to more favorable clinical outcomes in cancer patients.

**Conclusions:** Overall, our studies support pharmacological inhibition of MYSM1 DUB catalytic function as a novel therapeutic strategy for MYC-driven lymphoma and potentially other cancers.

## BACKGROUND

MYC is an oncogenic transcription factor, over-expressed, amplified, or otherwise dysregulated in over 50% of all cancers (1), including ∼90% of Burkitt lymphomas and ∼10% of diffuse large B-cell lymphomas (DLBCL) (2). Particularly in DLBCL, *MYC*-locus translocations are linked to rapid disease progression, poor therapy response, and inferior outcomes for the patients (2, 3, 4). Oncogenic activity of MYC is closely linked to its primary function as a transcriptional regulator of the genes engaged in ribosome biogenesis, protein synthesis, and cell growth (5, 6). In cancers with *MYC* chromosomal aberrations, sustained MYC activity is essential for tumor persistence, highlighting MYC as a critical therapeutic target (3, 7). However, direct targeting of MYC remains highly challenging due to its lack of ligand binding or catalytic domains and its intrinsically disordered structure. Drugs in development therefore aim to indirectly modulate MYC activity by targeting MYC-interacting proteins, antagonizing *MYC* gene expression, disrupting MYC dimerization with MAX, inhibiting their DNA-binding, or promoting degradation (8, 9). However, few of these compounds reached clinical trials, and new therapeutic approaches for MYC-driven cancers are urgently needed (10).

Myb-like SWIRM and MPN domains 1 (MYSM1) is a chromatin associated deubiquitinase (DUB) that regulates gene expression in hematopoietic and immune cells by removing ubiquitin from histone H2AK119, a repressive epigenetic mark, as well as other substrates (11). Recently, we and others demonstrated that MYSM1 maintains the normal expression of genes encoding ribosomal proteins and translation factors in non-malignant hematopoietic cells, and this activity is well conserved between mouse and human (12, 13). Cooperation of MYSM1 with MYC in the regulation of these transcriptional programs was supported by the strong overlap in their genomic binding particularly at these gene promoters (14). Building on this, we showed that a genetic ablation of *Mysm1*-expression offers protection against B cell lymphoma in the *EμMYC* murine model (14). However, major knowledge gaps remain – it is unclear whether the loss of MYSM1 DUB catalytic activity alone is sufficient for disease protection, whether this protective effect is conserved between mouse and human, and whether the loss of MYSM1 function in tumor cells, immune cells, and/or the tumor microenvironment plays a central role in the disease protection.

With the wide interest in DUBs as drug-targets (15, 16), we propose MYSM1 inhibition as a potential novel therapeutic strategy for MYC-driven B cell lymphoma. However, as some MYSM1 functions may be mediated via its many protein-protein interactions (11), it is now critical to assess whether a selective loss of MYSM1 DUB catalytic activity is sufficient to protect against lymphoma, without the loss of MYSM1 protein expression. To address this, we have established a novel mouse strain expressing MYSM1^D660N^ – with a mutation at the catalytic site that renders the protein fully inactive, but without affecting its normal expression and steady-state levels (17). Here, we apply this new mouse strain *Mysm1*^DN^ to assess whether the loss of MYSM1 DUB catalytic function alone can protect against lymphoma in the *EμMYC* model and to investigate the underlying mechanisms.

Antitumor immunity plays a central role in the control of cancer progression, and immunotherapies are revolutionizing cancer treatment in the clinic (18). Therefore, it is critical to understand how therapeutics developed to target the tumor also impact antitumor immunity. MYC is known to promote cancer immune evasion, by altering the tumor cell secretome and by inducing checkpoint marker expression (19, 20, 21). At the same time, MYC-regulated transcriptional programs in T cells are essential for their activation, clonal expansion, and the effective induction of antitumor immunity (22, 23, 24). MYSM1 is also widely investigated as a regulator of immune cell development and function (11) – in myeloid leukocytes of the innate immune system (25, 26, 27), in dendritic cells that bridge innate and adaptive immune responses (28, 29), and in lymphocytes and adaptive immunity (30, 31, 32). Nevertheless, the role of MYSM1 in antitumor immunity remains unexplored, and this represents an important knowledge gap towards establishing MYSM1 as a cancer drug-target.

In the current study, we for the first time test whether the loss of MYSM1 DUB catalytic activity, without the loss of its protein expression, is sufficient to protect against MYC-driven lymphoma in mouse models. We address the impact of MYSM1 on antitumor immunity – systematically interrogating how its function in the tumor cells, in different immune cell lineages, and in the tumor microenvironment (TME) impact the disease progression and the immune landscape of the tumors. Finally, leveraging the Cancer Dependency Map Project (DepMap) and Cancer Genome Atlas (TCGA) databases, we provide first evidence linking MYSM1 loss to impaired fitness of human lymphoma cell lines in culture and to favorable clinical outcomes in cancer patients.

## METHODS

### Mouse strains

Mouse strain B6.Cg-Tg(IghMyc)22Bri/J (JAX: 002728, abbreviated as *EμMYC*) is a widely used model of B cell lymphoma and overexpresses MYC under the control of the immunoglobulin heavy chain locus enhancer (33). Mouse strains *Mysm1*^fl/fl^, *Mysm1*^fl/fl^ Cre^ERT2^, and *EμMYC Mysm1*^fl/fl^ Cre^ERT2^ for a tamoxifen-induced *Mysm1*-gene deletion were previously described (12, 14, 34). Mouse strain *Mysm1*^D660N^ was established by Dr. Linda Henneman and Dr. Ivo J. Huijbers at the Netherlands Cancer Institute, Antoni Van Leeuwenhoek Ziekenhuis (Amsterdam, Netherlands), and carries a point mutation in the endogenous *Mysm1* gene to render the protein catalytically inactive, as we described previously (17). Mouse strains B6.129P2-*Lyz2*^tm1(cre)Ifo/J^ (JAX: 004781, LysM-Cre) and B6.Cg-Tg(Itgax-cre)-Reiz/J (JAX: 008068, CD11c-Cre) for gene deletion in myeloid cells and dendritic cells were purchased from the Jackson Laboratory (35, 36, 37). Mouse strains B6.Cg-Tg(Cd4-cre)1Cwi/BfluJ (JAX: 022071, CD4-cre) and B6.Cg-Tg(Lck-cre)548Jxm/J (JAX: 003802, Lck-cre) for gene deletion in T lymphocytes were purchased from Jackson laboratory (38, 39, 40). Note that the efficient deletion of *Mysm1*^fl/fl^ alleles in appropriate immune cell lineages in vivo with these Cre transgenes has been successfully validated in our previous work (28, 31). All the strains were on the C57BL/6 genetic background. The mice were maintained under specific pathogen-free conditions and sex-matched across the experimental groups. All experiments were in accordance with the guidelines of the Canadian Council on Animal Care and protocol MCGL-7643 approved by the McGill Animal Care Committee.

### Mouse genotyping

Mouse genotyping was performed with DreamTaq DNA Polymerase (ThermoFisher Scientific) or Kapa Taq DNA Polymerase (Millipore-Sigma) and primers from Integrated DNA Technologies, as described previously (14, 34). Genotyping of the *Mysm1*^D660N^ strain was performed with a custom designed TaqMan SNP Genotyping assay and the TaqMan Genotyping Master Mix on a StepOnePlus instrument (all from ThermoFisher Scientific), as described previously (17).

### Tamoxifen mouse treatment

Tamoxifen-induced *Mysm1*-gene deletion was carried out as previously described (14, 34), with minor modifications. Mice were injected intraperitoneally with tamoxifen (Millipore-Sigma, T5648) in sterilized corn oil at 0.12 mg/gram per injection, with 8 doses administered in total over 16 days. Successful deletion of *Mysm1* exon 3 was validated by PCR analyses of the genomic DNA from hematopoietic and lymphoid organs of the mice, as previously (34).

### Adoptive transfers of E***_μ_***MYC lymphoma cells

*EμMYC* tumor cells harvested from mice across *Mysm1* genotypes were processed to single cell suspensions, subjected to red blood cell lysis in ACK buffer (0.15M NH_4_Cl, 10mM KHCO_3_, 0.1mM EDTA) and cryopreserved in media consisting of 10% dimethyl sulfoxide (DMSO, Millipore-Sigma, D8418), 20% fetal calf serum (FCS, Thermo Fisher Scientific), and 70% B cell lymphoma culture media, corresponding to 45% IMDM (Thermo Fisher Scientific), 45% DMEM (Thermo Fisher Scientific), 10% FCS (Thermo Fisher Scientific), 100U/mL penicillin and 100µg/mL streptomycin (Wisent). Recipient mice were pre-conditioned with 1.75Gy or 3.5Gy whole body irradiation in an RS-2000 irradiator (Rad Source) and administered with tumor cells via intravenous injections at 1 million cells per mouse. Subsequently, the recipients were administered with tamoxifen or vehicle corn oil, as described above, and mouse health monitored. Survival was defined as the time to the terminal stage of disease, at which point the animals were euthanized according to protocol MCGL-7643 approved by the McGill University Animal Care Committee, and tumors were harvested for analyses.

### Cell culture

*EμMYC* tumor cells harvested and cryopreserved from the lymph nodes of *EμMYC* mice across the *Mysm1* genotypes were cultured on a monolayer of irradiated *Ink4a*^-/-^ mouse embryonic fibroblasts (MEFs) in media containing 45% DMEM (Thermo Fisher Scientific), 45% IMEM (Thermo Fisher Scientific), 10% FCS (Thermo Fisher Scientific), 100μg/mL streptomycin, 100U/mL penicillin (Wisent), and 5x10^-5^ M β-mercaptoethanol (Millipore-Sigma) (14).

### Flow cytometry

Cells were stained for surface markers in PBS with 2% fetal calf serum (Thermo Fisher Scientific) and 5% Super Bright Complete Staining Buffer (SB-4401-75, Thermo Fisher Scientific) for 20 minutes on ice with fluorophore conjugated antibodies, acquired from BioLegend, Thermo Fisher Scientific, or BD Biosciences, as listed in Supplemental Table S1. For intracellular staining the cells were fixed and permeabilized using the FOXP3 Transcription Factor Staining Buffer Set (00-5523-00, Thermo Fisher Scientific). The cells were stained with anti-eEF1G (EPR7200, Abcam) followed by Alexa Fluor 488 anti-rabbit IgG highly cross-adsorbed secondary antibody (Thermo Fisher Scientific), or appropriate isotype controls. For the analysis of intracellular reactive oxygen species (ROS), the CellROX Deep Red reagent was used according to the manufacturer’s protocol (C10422, Thermo Fisher Scientific). Fixable Viability Dye eFluor506 (65-0866-18, Thermo Fisher Scientific) was used to gate-out dead cells, and compensation was performed with BD CompBeads (552845, BD Biosciences) or UltraComp eBeads Plus (01-3333-42, Thermo Fisher Scientific). All data were acquired on BD Fortessa (BD Biosciences) and analyzed with FlowJo (BD Biosciences).

### Protein synthesis rate analysis

Analysis of protein synthesis was performed with the O-propargyl-puromycin (OPP) incorporation method. Cells were cultured in the presence of 20 μM OPP for 30 minutes, stained with Fixable Viability Dye eFluor506 (65-0866-18, Thermo Fisher Scientific), fixed in 2% paraformaldehyde in PBS with 2% FCS at 37 C for 10 minutes, and permeabilized in 90% methanol for 30 minutes on ice. Cells were washed in PBS and stained for OPP incorporation using the Click-iT Plus OPP Alexa Fluor 488 Protein Synthesis Assay kit (C10456, Thermo Fisher Scientific). Data were acquired on BD Fortessa (BD Biosciences) and analyzed with FlowJo (BD Biosciences).

### Fluorescence activated cell sorting and RNA isolation

For fluorescence activated cell sorting (FACS), cryopreserved cells were thawed in a 37°C water-bath and stained with fluorophore-conjugated antibodies: anti-IgM PE (ThermoFisher #12-5790-83), anti-B220-PerCP-Cy5.5 (BioLegend, #103236) and with the viability dye DAPI (BioLegend, #422801). Live, B220^+^IgM^−^ or B220^+^IgM^+^ cells were sorted on the FACSAria Fusion (BD Biosciences) into the Mag-MAX Lysis Buffer (Ambion, AM1830). RNA was isolated using the Mag-MAX total RNA kit (Ambion, AM1830) according to the manufacturer’s protocol, with RNaseOUT Recombinant Ribonuclease Inhibitor (Invitrogen, 10777019) added to buffers. RNA yields and quality were assessed on Bioanalyzer (Agilent).

### RNA sequencing

Bulk RNA sequencing was performed as previously described with minor modifications (12, 41). In brief, total RNA was extracted using the MagMAX total RNA kit (Ambion), and purification was carried out with RNAClean XP (Beckman Coulter). RNA integrity was evaluated using Bioanalyzer RNA Pico chips (Agilent). rRNA depletion and library construction were completed using the KAPA RiboErase and RNA HyperPrep Kit (Roche Diagnostics). Libraries were sequenced on an Illumina NovaSeq PE159 platform in paired-end 100 bp mode, targeting approximately 50×10 reads per sample. Read quality was assessed with the FastQC tool (Babraham Bioinformatics), and low-quality bases were trimmed from read ends using Trimmomatic v.0.33 (42). Sequencing reads were aligned to the mouse UCSC GRCm38/mm10 reference genome using Hisat2 v2.2.1 (43, 44, 45). Gene expression was quantified based on uniquely mapped reads using featureCounts with default parameters (Subread package v1.5.2) (46). Residual rRNA reads were removed, and only genes expressed above 5 counts per million reads (CPM) in at least 3 samples were retained, for a total of 37,716 genes. Dimension reduction analysis was performed using the Principal Component Analysis in R (prcomp function). Normalization and differential expression analysis were carried out using the edgeR Bioconductor package (47). Genes showing ≥ |1.5|-fold change with a Benjamini–Hochberg adjusted p value ≤ 0.001 were considered significantly differentially expressed. Gene Ontology (GO) enrichment analysis of differentially expressed genes was performed using DAVID Bioinformatics Resources 6.8 within the Biological Processes (BP4) category (48, 49). Gene Set Enrichment Analysis (GSEA) was conducted with GSEA v4.3.2 using MSigDB v2025.1 under default parameters and gene set–based permutation (50). *De novo* motif finding analyses were done using Hypergeometric Optimization of Motif EnRichment (HOMER v5.1, 7-16-2024) (51). RNA-Seq datasets from our study have been deposited into the NCBI public database and are available under the GEO accession number GSE310117.

### DepMap and cBioPortal cancer genomics databases

To assess the impact of MYSM1 loss-of-function on the fitness of human lymphoma cells, the Cancer Dependency Map Project database was interrogated for CRISPR-screen data for *MYSM1* gene and lymphoid malignancies (DepMap, https://depmap.org, 23Q4) (52). Chronos dependency scores were then compared either between cell lines or to zero, which corresponds to the null hypothesis of ‘no effect of MYSM1 loss on cell fitness’. Information on the age and sex of the patients that the cell lines were derived from, as well as on the *Tp53* status of the cells and the presence of *MYC*, *BCL2*, and *BCL6* locus translocations were obtained manually for each line from the databases of DSMZ-German Collection of Microorganisms and Cell Cultures (www.dsmz.de) or from the American Type Culture Collection (ATCC, www.atcc.org).

The Cancer Genome Atlas (TCGA) – PanCancer Atlas was interrogated via cBioPortal for data on *MYSM1* somatic mutation, copy number variation, and patient clinical data across 32 non-redundant studies spanning many cancer types (www.cbioportal.org) (53, 54, 55, 56). Somatic mutation data were filtered to include only non-silent *MYSM1* mutations, defined as missense, nonsense, frameshift, and splice-site variants. Patients without *MYSM1* mutations were classified as wild type. For copy number, gain represents a low-level gain of copy number whereas amplification indicates a higher-level gain of copy number, derived from copy-number analysis algorithms like GISTIC(57) or RAE(58). Clinical outcome data, including overall survival, disease-specific survival, disease-free interval (DFI), and progression-free interval (PFI), were extracted for matched samples. Kaplan–Meier survival analyses were performed to compare outcomes between *MYSM1* mutant and *MYSM1* wild-type groups, using the log-rank test (Mantel-Cox method) and Gehan-Breslow-Wilcoxon test in Prism 10.2.2 (GraphPad Inc).

### Statistical analyses

Statistical comparisons were performed with Prism 9.5.1-10.5.0 (GraphPad Inc.), using Student’s *t-*test for two groups, ANOVA for multiple comparisons, and Kaplan–Meier regression analysis for survival data.

## RESULTS

### Loss of MYSM1 catalytic activity is sufficient to protect against *E***_μ_***MYC* lymphoma

To test the impact of a selective loss of MYSM1 DUB catalytic activity on lymphoma, the lymphoma-prone *EμMYC* mice (33) were crossed to the *Mysm1*^DN^ mouse strain that expresses a catalytically inactive MYSM1^D660N^ (17). Survival of the *EμMYC Mysm1*^DN/DN^ mice was compared to *EμMYC* littermates of control *Mysm1* genotypes. This demonstrated highly significant extension in the survival of the *EμMYC Mysm1*^DN/DN^ mice relative to control littermates, establishing that the loss of MYSM1 catalytic activity can protect against MYC-driven lymphoma (Figure 1A).

**Figure 1.**
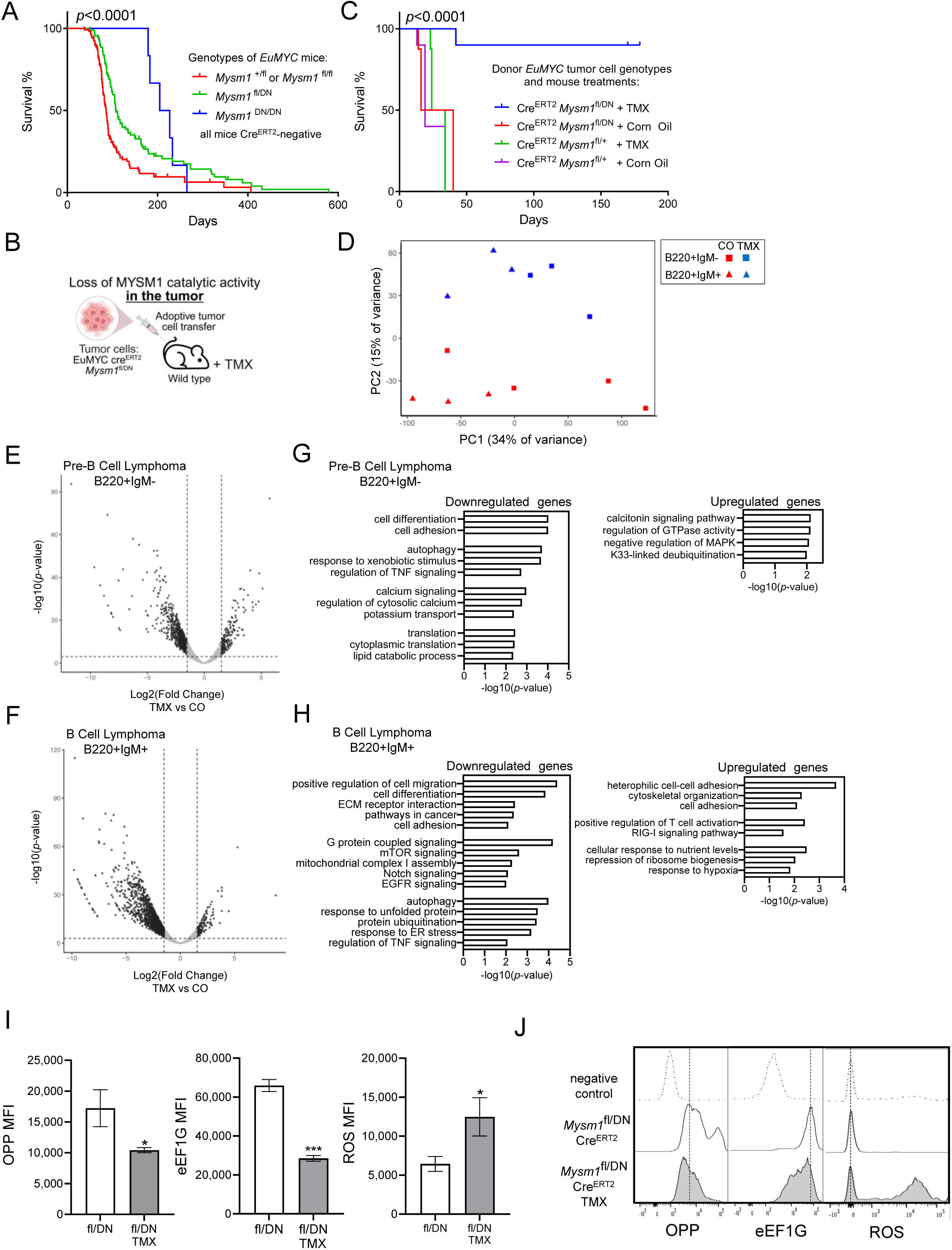
Loss of MYSM1 DUB catalytic activity protects against B cell lymphoma onset and progression, via its impact on the cancer cell transcriptome and physiology. (A) Survival of *EμMYC Mysm1*^DN/DN^ mice (n=6) relative to littermates of control genotypes, including *EμMYC Mysm1*^fl*/*DN^ (n=92), and *EμMYC Mysm1*^fl/+^ or *EμMYC Mysm1*^fl/fl^ (n=108). Note that in the absence of Cre-transgene and tamoxifen induction the *Mysm1*^fl^ allele expresses wild type MYSM1. *p*-values were calculated using log-rank Mantel-Cox test. (B) Schematic of the tumor adoptive cell transfer model, with wild type mice receiving *EμMYC* Cre^ERT2^ lymphoma cells of *Mysm1*^fl/DN^ or control *Mysm1*^fl/+^ genotypes followed by either tamoxifen (TMX) for Cre^ERT2^-mediated *Mysm1*^fl^ allele deletion or vehicle corn oil. (C) Survival of wild-type mice after an adoptive transfer of 1 million *EμMYC* Cre^ERT2^ tumor cells of *Mysm1*^fl/DN^ or control *Mysm1*^fl/+^ genotypes, followed with either TMX or control corn oil treatment – loss of MYSM1 catalytic activity protects against lymphoma progression. *p*-values calculated using log-rank Mantel-Cox test with n=8-10 mice per group; conclusions reproduced in a second independent experiment. (D-H) Loss of MYSM1 catalytic activity alters the tumor cell transcriptome, as shown with bulk RNA-sequencing analyses of *EμMYC* Cre^ERT2^ lymphoma cells comparing between *Mysm1*^Δ/DN^ and control *Mysm1*^fl/DN^ genotypes. (D) Principal component analysis of the gene expression profiles of *EμMYC* tumors, with PC1 providing some separation between the B220^+^IgM^−^ pre-B and B220^+^IgM^+^ mature B cell lymphomas (34% of variance) and PC2 clearly separating *Mysm1*^Δ/DN^ from control *Mysm1*^fl/DN^ samples (15% of variance). (E-F) Volcano plots showing the transcriptional profiles of *EμMYC* Cre^ERT2^ lymphomas of *Mysm1*^Δ/DN^ versus control *Mysm1*^fl/DN^ genotypes, including B220^+^IgM^−^ pre-B and B220^+^IgM^+^ mature lymphomas. Differentially expressed genes are defined based on fold change (FC) ≥ |1.5| and false discovery rate (FDR) ≤ 0.001; full gene lists are provided in Table S2. (G-H) Gene ontology (GO) analysis showing the enriched biological processes in the bulk RNA-seq transcriptional data of *EμMYC* Cre^ERT2^ lymphomas of *Mysm1*^Δ/DN^ versus control *Mysm1*^fl/DN^ genotypes, including B220^+^IgM^−^pre-B and B220^+^IgM^+^ mature lymphomas. Dysregulated genes are defined at FC ≥ |1.5| and FDR ≤ 0.001; -log10(*p*-values) for select top GO-terms are graphed and the full GO-term analyses results are provided in Table S3. (I-J) Loss of MYSM1 catalytic activity in tumor cells results in reduced protein synthesis, reduced levels of translation factor eEF1G, and increased levels of reactive oxygen species (ROS). *EμMYC* Cre^ERT2^ *Mysm1*^fl/DN^ tumor cells from TMX-induced or control corn oil treated mice were analyzed in culture for protein synthesis (OPP assay), eEF1G translation factor, and ROS levels through flow cytometry. Mean fluorescence intensity (MFI) is plotted, and representative flow cytometry plots showing OPP incorporation, eEF1G staining, and ROS levels are presented. Controls represent non-specific staining of cells not exposed to OPP, cells stained with an isotype control antibody, or cells not loaded with the ROS-sensitive CellROX dye, respectively. Bars represent mean ± SEM from n=3 tumor bearing mice per genotype. Statistical analyses used Student’s *t*-test; * *p*<0.05, *** *p*<0.001.

To assess the role of MYSM1 DUB catalytic activity specifically in disease progression, we crossed *Mysm1*^DN^ mice to the *EμMYC* Cre^ERT2^ *Mysm1*^fl/fl^ mouse strain that allows an inducible deletion of *Mysm1*^fl^ allele with tamoxifen administration (14, 34). Adoptive tumor cell transfer model was applied to test the effects of the loss of MYSM1 catalytic activity on lymphoma progression, independently of developmental phenotypes, rates of disease onset, or of MYSM1 functions in the tumor microenvironment. *EμMYC* Cre^ERT2^ tumors of *Mysm1*^fl/DN^ or control *Mysm1*^fl/+^ genotypes were harvested from terminally ill donor mice and tumor cells adoptively transferred into wild type recipient mice, at 1 million cells per mouse. Cohorts of recipients subsequently received either tamoxifen for Cre^ERT2^-induced *Mysm1*^fl^ to *Mysm1*^Δ^ allele conversion or control vehicle corn-oil (Figure 1B). The study demonstrated that the loss of MYSM1 catalytic activity in tumor cells, corresponding to the mouse cohort receiving *EμMYC* Cre^ERT2^ *Mysm1*^fl/DN^ tumors followed by tamoxifen, resulted in a highly significant delay in disease progression and extension in mouse survival (Figure 1C). We conclude that the loss of MYSM1 catalytic activity is sufficient to protect against lymphoma progression, establishing MYSM1 as a novel drug-target.

To analyze the tumor-intrinsic mechanisms that lead to the protection against disease with the loss of MYSM1 catalytic activity, bulk RNA-seq transcriptional analyses were performed on *EμMYC* Cre^ERT2^ lymphoma cells of *Mysm1*^Δ/DN^ and control *Mysm1*^fl/DN^ genotypes, using FACS-sorted live B220^+^IgM^−^ pre-B and B220^+^IgM^+^ mature lymphoma cells from primary tumors. Principal component analysis revealed some segregation of the gene expression profiles of pre-B and mature B cell lymphomas (PC1, 34% of variance), and a clear segregation of the gene expression based on the *Mysm1* genotype (PC2, 15% of variance, Figure 1D). Differential gene expression analysis compared *EμMYC* Cre^ERT2^ lymphoma cells of *Mysm1*^Δ/DN^ versus control *Mysm1*^fl/DN^ genotypes, at fold change (FC) ≥ |1.5| and false discovery rate (FDR) ≤ 0.001. This identified 815 downregulated and 157 upregulated genes in pre-B cell lymphomas (Figure 1E, Tables S2A-B), and 1323 downregulated and 137 upregulated genes in mature B cell lymphomas (Figure 1F, Tables S2C-D). This for the first time demonstrates the impact of the loss of MYSM1 function on the tumor cell transcriptome, and is consistent with the established MYSM1 role as an activator of gene expression. Gene ontology (GO) analyses demonstrated dysregulation of the transcriptional programs linked to cell adhesion, migration, as well as various signaling and stress response pathways with the loss of MYSM1 function (Figures 1G-H, Table S3). Importantly, there was also a downregulation of the transcriptional signatures of protein translation and mTOR signaling, and an upregulation of immunological transcriptional signatures (Figures 1G-H, Table S3). Further gene set enrichment (GSEA) analyses supported the downregulation of transcriptional signatures linked to translation and protein synthesis particularly in mature lymphoma cells, and the upregulation of immunological transcriptional signatures across both pre-B and mature B cell lymphomas (Figure S1A-B, Table S4). This is consistent with previously reported role of MYSM1 in normal blood and immune cells to sustain the gene expression programs of ribosome biogenesis and protein translation (12, 13, 14). It also suggests that the protection against lymphoma disease with the loss of MYSM1 catalytic function may arise via both an intrinsic disruption of tumor cell physiology and its potential impact on antitumor immunity.

*EμMYC* lymphoma cells across the *Mysm1* genotypes were further analyzed in culture, to reveal the mechanisms protecting against lymphoma progression with the loss of MYSM1 catalytic activity. *EμMYC Mysm1*^Δ/DN^ lymphoma cells had significantly lower protein synthesis rates and levels of the eEF1G translation factor, and significantly higher reactive oxygen species levels than control *EμMYC Mysm1*^fl/DN^ lymphoma cells (Figure 1I-J). This is consistent with previously established models describing the co-regulation of genes encoding ribosomal and translation machinery by MYSM1 and MYC in hematopoietic cells (12, 13), establishing the mechanisms through which tumor-intrinsic loss of MYSM1 protects against lymphoma.

### Loss of MYSM1 catalytic activity in tumor cells indirectly modulates antitumor immunity

To explore whether the loss of MYSM1 catalytic activity in the tumor cells may also indirectly modulate antitumor immunity, we analyzed lymphoma cells of test and control *Mysm1* genotypes for expression of co-stimulatory molecules CD86, CD80, CD40, as well as MHCII and checkpoint marker PD-L1. Notably, with the loss of MYSM1 activity *EμMYC Mysm1*^Δ/DN^ tumor cells showed a moderate upregulation of CD80 and PD-L1 relative to control *EμMYC Mysm1*^fl/DN^ tumor cells, but no significant changes in the other markers (Figure S1C-D). Overall, this suggested that loss of MYSM1 catalytic activity within the tumor may modulate antitumor immunity.

To further explore how the loss of MYSM1 catalytic activity in tumor cells affects antitumor immunity, the *EμMYC* Cre^ERT2^ tumors of *Mysm1*^fl/DN^ or control *Mysm1*^fl/+^ genotypes harvested from tamoxifen treated or control corn oil treated mice were analyzed for immune cell infiltration. We observed a highly significant increase in the abundance of dendritic cells (DCs), including both the conventional (cDC1, cDC2) and plasmacytoid (pDC) subsets, in the *Mysm1*^Δ/DN^ relative to control tumors (Figure 2A-B). No changes in cDC activation were observed between the groups, as assessed by CD86, CD80, CD40, MHCII, and PD-L1 expression (Figure S2A-B).

**Figure 2.**
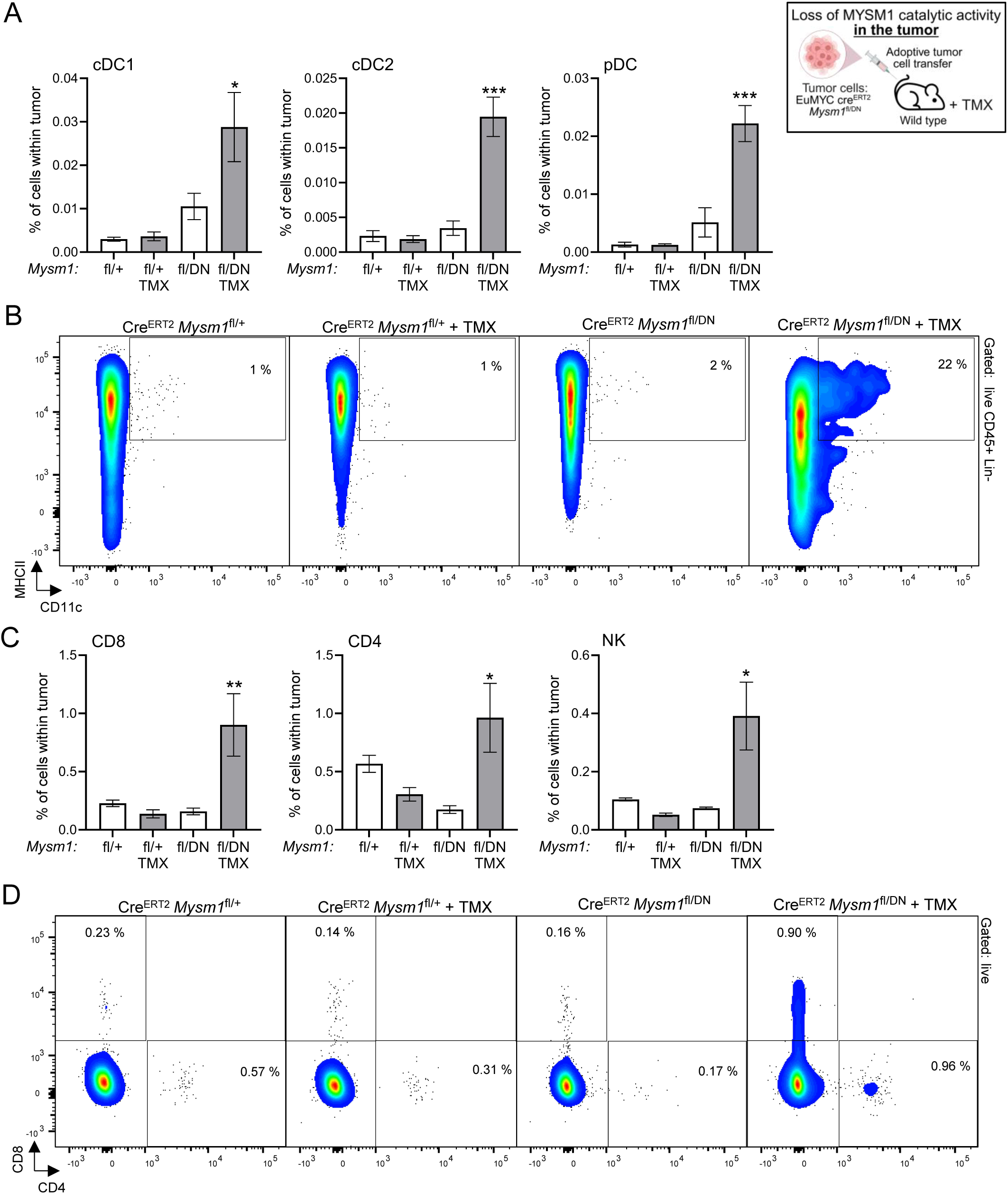
Loss of MYSM1 catalytic activity in the tumor cells results in enhanced immune cell infiltration into the tumor. (A-B) Increased infiltration of dendritic cells into the tumors following an inducible tumor-intrinsic loss of MYSM1 DUB catalytic activity. **(A)** Frequencies of cDC1 and cDC2 conventional dendritic cells and pDC plasmacytoid dendritic cells in the tumors of Cre^ERT2^ *Mysm1*^fl/+^ and Cre^ERT2^ *Mysm1*^fl/DN^ genotypes, following the treatment with tamoxifen (TMX) to induce Cre^ERT2^ activation or with corn oil as the vehicle control. Bars represent mean ± SEM from n=4-10 tumor bearing mice per genotype, consolidated from two independent experiments. Statistical analyses used ANOVA with Sidak’s posthoc test, comparing the tumors from TMX-induced and control mice, * *p*<0.05, *** *p*<0.001, and not significant if not indicated. DCs were gated as live CD45^+^Lin^−^F4/80^−^CD64^−^CD11c^+^MHCII^+^ cells, followed by B220^−^PDCA1^−^XCR1^+^SIRPα^−^ for cDC1, B220^−^PDCA1^−^XCR1^−^SIRPα^+^ for cDC2, and B220^+^ PDCA1^+^ for pDCs, with the gating presented in Figure S7. **(B)** Representative flow cytometry plots of the tumor samples gated on live CD45^+^Lin^−^F4/80^−^CD64^−^ cells and showing the CD11c^+^MHCII^+^ dendritic cell gate. Numbers on the plots indicate the percentages of cells within each gate out of the parent population, presented as mean for all the tumor samples of that genotype and treatment group. **(C-D)** Increased infiltration of T cells and NK cells into the tumors following an inducible tumor-intrinsic loss of MYSM1 DUB catalytic activity. **(C)** Frequencies of CD8 T cells, CD4 T cells, and NK cells in the tumors of Cre^ERT2^ *Mysm1*^fl/+^ and Cre^ERT2^ *Mysm1*^fl/DN^ genotypes, following the treatment with TMX to induce Cre^ERT2^ activation or with corn oil as the vehicle control. Bars represent mean ± SEM from n=4-10 tumor bearing mice per genotype, consolidated from two independent experiments. Statistical analyses used ANOVA with Sidak’s posthoc test, comparing the tumors from tamoxifen treated and control mice; * *p*<0.05, ** *p*<0.01, and not significant if not indicated. T cells were gated as live B220^−^CD3^+^CD8^+^CD4^−^ and B220^−^CD3^+^CD4^+^CD8^−^ cells, and NK cells as live B220^−^CD3^−^NK1.1^+^. **(D)** Representative flow cytometry plots of the tumor samples gated on live cells and showing CD8 and CD4 T cells. Numbers on the plots indicate the percentages of cells within each gate out of the parent population, presented as the mean for all the tumor samples of that genotype and treatment group.

*Mysm1*^Δ/DN^ tumors also showed strongly increased infiltration of CD8 T cells, CD4 T cells, and NK cells relative to control tumors (Figure 2C-D). There were no significant differences in CD4 or CD8 T cell activation or exhaustion between the groups, as assessed by CD44, CTLA4, PD1, LAG3, and TIM3 markers, except for some upregulation of PD1 on CD8 T cells (Figure S2C & data not shown). There were also no differences in the proportions of tumor infiltrating CD4 T cells positive for transcription factors Tbet, GATA3, RORγT, or FOXP3, corresponding to Th1, Th2, Th17, and Treg lineages, between *Mysm1*^Δ/DN^ and control tumors (Figure S2D-F). Distribution of tumor infiltrating CD4 or CD8 T cells between the CD62L^+^CD44^−^ naive and CD62L^−^CD44^+^ antigen experienced subsets was also unchanged (Figure S2G).

Analyses of myeloid cells in *Mysm1*^Δ/DN^ relative to the control tumors demonstrated an increased abundance of monocyte derived macrophages, with no changes in neutrophils or tumor resident macrophages (Figure S2H-I). Assessment of myeloid cell activation based on CD86, CD80, CD40, MHCII and PD-L1 demonstrated a significant increase in the activation marker CD86 on tissue resident macrophages in the *Mysm1*^Δ/DN^ relative to control tumors, with no differences for other markers and cell types (Figure S2J-K & data not shown). Flow cytometry gating strategies to characterize tumor infiltrating immune cells are presented in Figures S7-9.

Overall, loss of MYSM1 catalytic activity in the *EμMYC* tumor cells resulted in significantly increased infiltration of DCs, T cells, and NK cells into the tumors. Given the major role of these leukocytes as mediators of antitumor immune response (59, 60, 61), this suggests that the loss of MYSM1 function within the tumor may act to enhance antitumor immunity.

### MYSM1 loss-of-function in immune cells: impact on survival and antitumor immunity

Our results above support the targeting of MYSM1 DUB catalytic activity as a novel therapeutic approach for MYC-driven lymphoma. However, MYSM1-regulated transcriptional programs have been widely implicated in immune cell development and function (11, 22, 23, 24). Therefore, the impact of targeting MYSM1 on antitumor immunity requires further investigation. We transferred wild type *EμMYC* lymphoma cells into cohorts of recipient mice of *Mysm1*^DN/fl^, *Mysm1*^fl/fl^ and control *Mysm1*^fl/+^ and *Mysm1*^+/+^ genotypes, transgenic for DC-specific CD11c-cre (35), myeloid lineage specific LysM-Cre (36, 37), and T cell specific CD4-cre and Lck-cre (38, 39, 40). Note that efficient deletion of *Mysm1*^fl/fl^ alleles in appropriate immune cell lineages in vivo with these Cre transgenes has been successfully validated in our previous studies (28, 31). The mice were analyzed for disease progression, survival, and for immune cell infiltration into the tumors of terminally ill animals, comparing across the *Mysm1* genotypes of the recipients.

DC-specific loss of *Mysm1* or its catalytic activity had no significant impact on lymphoma progression and mouse survival (Figure 3A, CD11c-cre). Nevertheless, it resulted in an enhanced infiltration of cDCs, CD8 T cells, CD4 T cells, and monocyte derived macrophages into the tumors (Figure 3B). Activation and exhaustion markers on tumor infiltrating DCs, including CD86, CD80, CD40, MHCII and PD-L1, were not significantly altered (Figure S3A). Notably, the proportion of Tbet-positive tumor infiltrating CD4 T cells was significantly increased, indicative of their Th1-polarization (Figure S3B), and there was a significant increase in the activation markers CD86 and CD80 on monocyte derived macrophages (Figure S3C). Other CD4 T-helper cell subsets measured by GATA3, RORγt, or FOXP3 expression, and the activation of tumor infiltrating monocytes and tissue resident macrophages were not significantly affected (data not shown). In summary, loss of MYSM1 or its DUB catalytic activity in DCs results in enhanced immune infiltration and activation within the tumors, but without a significant impact on disease progression and survival.

**Figure 3.**
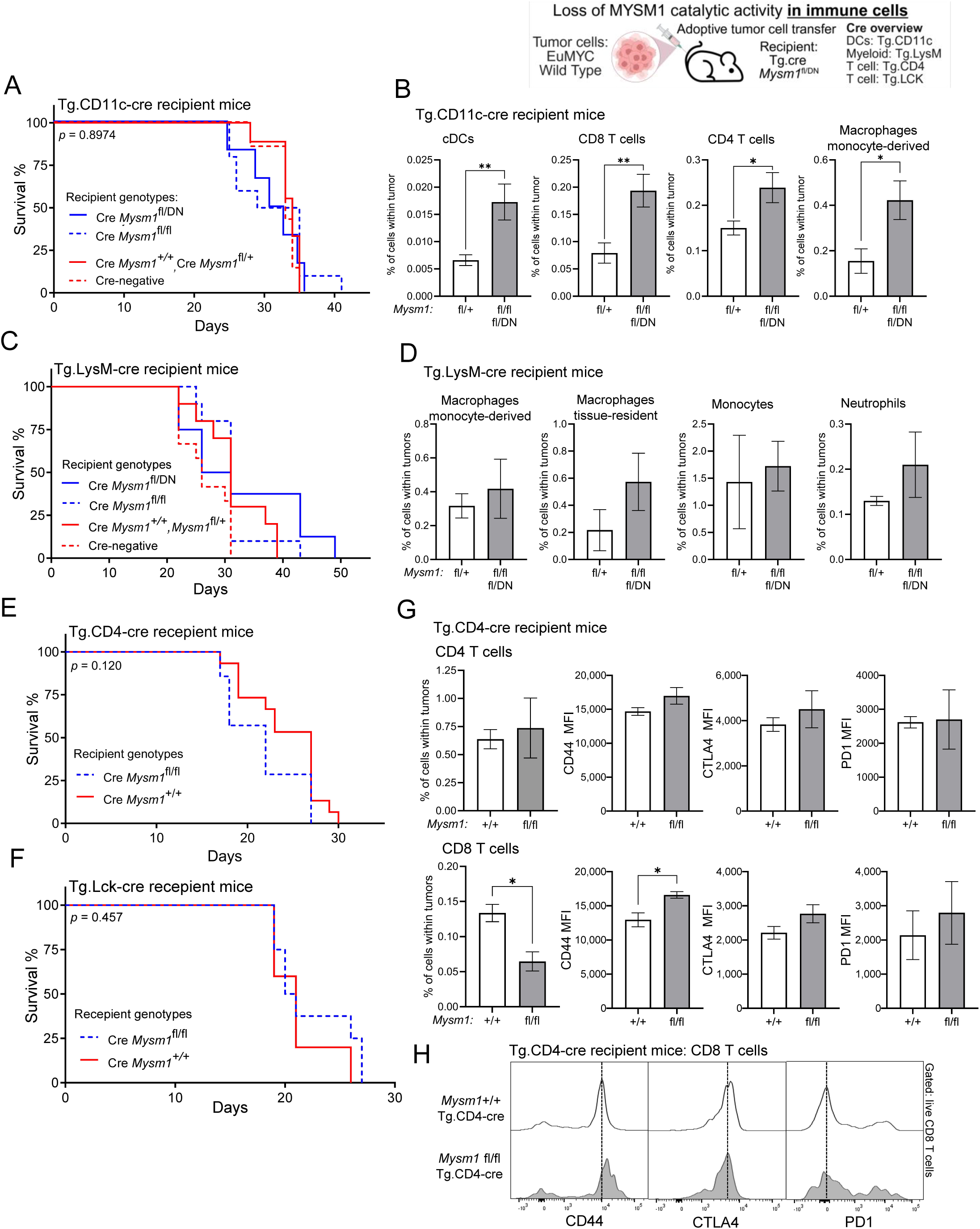
Impact of the loss of MYSM1 function in immune cells on lymphoma progression, mouse survival, and immune cell infiltration into the tumors. Wild type *EμMYC* lymphoma cells were adoptively transferred into Cre transgenic recipient mice of *Mysm1*^DN/fl^, *Mysm1*^fl/fl^ and control *Mysm1*^fl/+^ and *Mysm1*^+/+^ genotypes, analyzing mouse survival and immune cell infiltration into the tumors of terminally ill mice. The following Cre-transgenic lines were used: CD11c-cre for *Mysm1* deletion in dendritic cells (DCs) (35), LysM-Cre for *Mysm1* deletion in myeloid lineage cells (36, 37), and CD4-cre and Lck-cre for *Mysm1* deletion in T cells (38, 39, 40). Conventional DCs were gated as live CD45^+^Lin^−^B220^−^F4/80^−^CD64^−^CD11c^+^MHCII^+^ cells; myeloid lineage cells were gated as live CD45^+^B220^−^ cells, followed by Ly6G^−^Ly6C^+^ F4/80^−^ for monocytes, Ly6G^−^Ly6C^+^F4/80^+^ for monocyte derived macrophages, Ly6G^−^Ly6C^−^F4/80^+^ for tissue resident macrophages, and Ly6G^+^ for neutrophils; T cells were gated as live CD45^+^B220^−^NK1.1^−^CD3^+^, followed by CD4^−^CD8^+^ or CD4^+^CD8^−^, as shown in Figures S7-9. **(A)** Loss of *Mysm1* or its catalytic activity in DCs has no significant impact on lymphoma progression and mouse survival; *p*-values calculated using log-rank Mantel-Cox test; n=6-10 mice per group. **(B)** Loss of *Mysm1* or its catalytic activity in DCs results in an enhanced infiltration of cDCs, CD8 T cells, CD4 T cells, and monocyte derived macrophages into the tumors. **(C)** Loss of *Mysm1* or its catalytic activity in myeloid lineage immune cells has no significant impact on lymphoma progression and mouse survival; *p*-values calculated using log-rank Mantel-Cox test; n=8-10 mice per group. **(D)** Loss of *Mysm1* or its catalytic activity in myeloid cells has no significant effect on the abundance of monocyte derived macrophages, tissue resident macrophages, monocytes, or neutrophils within the tumors. **(E-F)** Loss of *Mysm1* in T lymphocytes has no significant effect on lymphoma progression and mouse survival; *p*-values calculated using log-rank Mantel-Cox test; n=7-15 mice per group for CD4-cre and n=5-8 mice per group for Lck-cre studies. **(G-H)** Loss of *Mysm1* in T lymphocytes results in reduced abundance of CD8 T cells within the tumors, and in an increased expression of CD44 activation marker on tumor infiltrating CD8 T cells, with no significant changes in the abundance of CD4 T cells, or in the expression of CTLA4 and PD1 exhaustion markers. Bars represent mean ± SEM; MFI – mean fluorescence intensity; statistical analyses used Student’s *t*-test, * *p*<0.05, ** *p*<0.01, or not significant if not indicated.

Loss of *Mysm1* or its DUB catalytic activity in myeloid lineage cells also had no significant impact on lymphoma progression and mouse survival (Figure 3C, LysM-cre). Furthermore, immune cell infiltration and activation within the tumors in this model were not significantly affected (Figure 3D, Figure S3D, data not shown). Finally, loss of *Mysm1* in T lymphocytes also had no significant impact on lymphoma progression and survival, in either CD4-cre or Lck-cre recipient models (Figure 3E-F). Analyzing immune cell infiltration and activation in the tumors in the CD4-cre recipient model, we find the reduced abundance of CD8 but not CD4 T cells (Figure 3G), as well as an increase in activation marker CD44 on tumor infiltrating CD8 T cells (Figure 3G-H). Other markers of T cell activation, polarization, and exhaustion were not significantly altered on either CD8 or CD4 T cells (Figure 3G-H, Figure S4A-C, data not shown), and there were no downstream effects on infiltration or activation of DCs or other myeloid lineage immune cells within the tumors (data not shown). Overall, we conclude that the selective loss of MYSM1 in immune cells did not impact lymphoma progression or mouse survival, but had significant modulatory effects on immune cell infiltration and activation within the tumors.

### MYSM1 loss in tumor microenvironment: impact on survival and antitumor immunity

To test if systemic loss of MYSM1 or its DUB catalytic activity in the tumor microenvironment (TME) can affect lymphoma progression, mouse survival, and antitumor immunity, wild type *EμMYC* lymphoma cells were adoptively transferred into Cre^ERT2^ transgenic recipients of *Mysm1*^fl/fl^, *Mysm1*^DN/fl^, and control *Mysm1*^fl/+^ genotypes. All the mice were administered with tamoxifen to induce Cre^ERT2^ activation in the TME (Figure 4A), with efficient deletion of *Mysm1*^fl^ allele in vivo in the Cre^ERT2^ model successfully validated in our previous studies (12, 17, 34).

**Figure 4.**
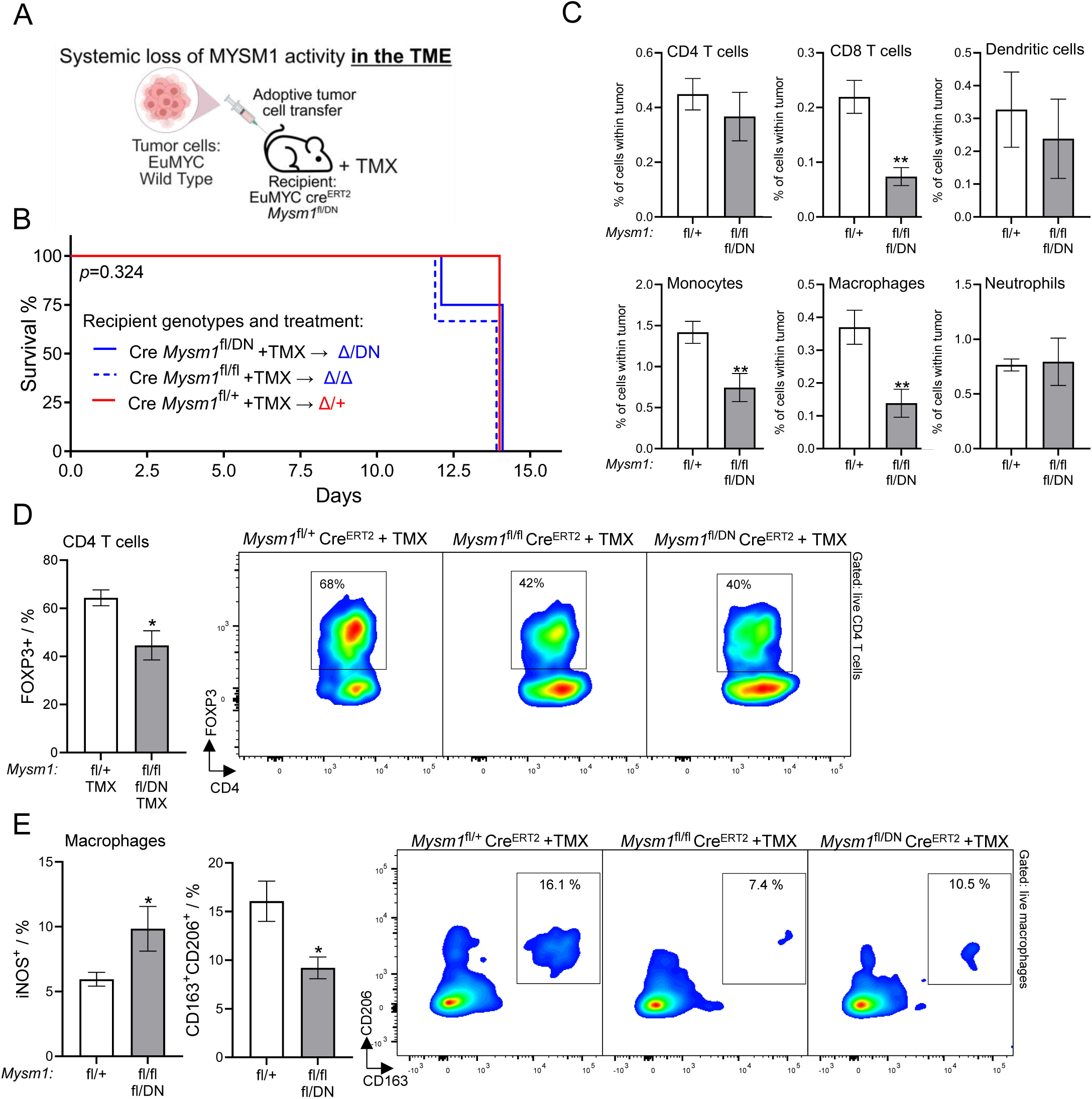
Impact of a systemic loss of MYSM1 function in the tumor microenvironment on lymphoma progression, mouse survival, and immune cell infiltration into the tumors. **(A)** Schematic summary of the experimental setup: wild type *EμMYC* lymphoma cells were adoptively transferred into Cre^ERT2^ transgenic recipient mice of *Mysm1*^DN/fl^, *Mysm1*^fl/fl^, and control *Mysm1*^fl/+^ genotypes and mouse survival compared following tamoxifen (TMX) treatment. **(B)** Impact of the loss of MYSM1 function in the tumor microenvironment (TME) on disease progression and mouse survival; *p*-values from log-rank Mantel-Cox test show no significant differences; n=3-7 mice per group. **(C-E)** Impact of the loss of MYSM1 function in the TME on immune cell infiltration. Bars represent mean ± SEM from n=7-8 mice per group; MFI – mean fluorescence intensity. Statistical analyses used Student’s *t*-test, * *p*<0.05, ** *p*<0.01, or not significant if not indicated. **(C)** Quantification of tumor infiltrating immune cells as a proportion of live cells within the tumor, including CD4 and CD8 T cells, dendritic cells, monocytes, macrophages, and neutrophils. T cells are gated as live CD45^+^B220^−^NK1.1^−^CD3^+^, followed by CD4^−^CD8^+^ or CD4^+^CD8^−^; DCs are gated as live CD45^+^Lin^−^F4/80^−^CD64^−^CD11c^+^MHCII^+^ cells; myeloid cells are gated as live CD45^+^B220^−^ followed by Ly6G^−^Ly6C^+^F4/80^−^ for monocytes, Ly6G^−^F4/80^+^ for macrophages, and Ly6G^+^ for neutrophils. **(D)** Impact of a systemic loss of MYSM1 function in the TME on tumor infiltrating CD4 T cells: proportion of tumor infiltrating CD4 T cells with positive intracellular staining for transcription factor FOXP3, and representative flow cytometry plots showing the gating on CD4^+^FOXP3^+^ tumor infiltrating Treg cells. **(E)** Impact of a systemic loss of MYSM1 function in the TME on tumor infiltrating macrophages: analyses for iNOS as a marker of classical (M1) activation and for CD163 and CD206 as markers of alternative (M2) activation; flow cytometry plots of the tumors showing a reduced proportion of macrophages expressing CD163 and CD206 markers of alternative (M2) activation following a systemic loss of MYSM1 function in the TME.

Systemic loss of MYSM1 or its catalytic activity in the TME had no significant effect on disease progression, with no differences in survival of *Mysm1*^Δ/Δ^, *Mysm1*^DN/Δ^, and control *Mysm1*^Δ/+^ tumor recipient mice (Figure 4B). Notably, *Mysm1*-deficient hosts showed a significant decrease in tumor infiltrating CD8 T cells, monocytes, and macrophages, while tumor infiltrating CD4 T cells, DCs, and neutrophils remained unchanged (Figure 4C). Nevertheless, loss of MYSM1 in the TME also resulted in a significant decrease in FOXP3^+^ Treg-like CD4 T cells (Figure 4D), an increase in iNOS+ M1-like classically activated macrophages, and a decrease in CD163+CD206+ M2-like alternatively activated macrophages (Figure 4E), all of which are expected to have a positive impact on antitumor immunity. Other markers of T cell activation, polarization, and exhaustion were not significantly affected (Figure S5A-F). There were some shifts in activation markers on tumor infiltrating monocytes, with an increase in CD86 and a reduction in CD80 and PD-L1, while the activation of tumor infiltrating DCs was unaffected (Figure S5G-J).

Overall, we conclude that a systemic loss of MYSM1 function in the TME has no significant impact on lymphoma progression and survival, at least within the timeframe of this experimental model (Figure 4A-B), but results in some reduction in immune cell infiltration into the tumors (Figure 4C). Systemic loss of MYSM1 in the TME also modulates the activation and polarization of tumor infiltrating immune cells, with reduction in FOXP3^+^ CD4 Tregs and enhanced M1 macrophage polarization (Figure 4D-E), which may have positive effects on antitumor immunity.

### Protective effects of MYSM1 loss in human lymphoma

To assess the impact of MYSM1 loss-of-function on the fitness of human lymphoma cells, we interrogated the Cancer Dependency Map Project (DepMap, depmap.org) (52) for CRISPR-knockout screen data for *MYSM1* gene in lymphoid malignancies. This demonstrated a highly significant loss of cell fitness with the loss of *MYSM1* for all human lymphoid cancer cell lines, as well as specifically for B cell lymphoma cell lines (Figure 5A-B). The protective effect of *MYSM1* loss was independent of the sex or age of the patient the tumor cells derived from (Figure S6A-B), and also independent of the *Tp53* status of the cells (Figure 5C). Notably, the double-hit or triple-hit lymphoma cell lines, characterized by *MYC*-locus translocations together with *BCL2* and/or *BCL6* translocations, were significantly more sensitive to *MYSM1* loss relative to other lymphoid malignancies (Figure 5D). Such cell lines represent the most aggressive lymphomas, prone to disease relapse and in need of novel treatment approaches (62, 63).

**Figure 5.**
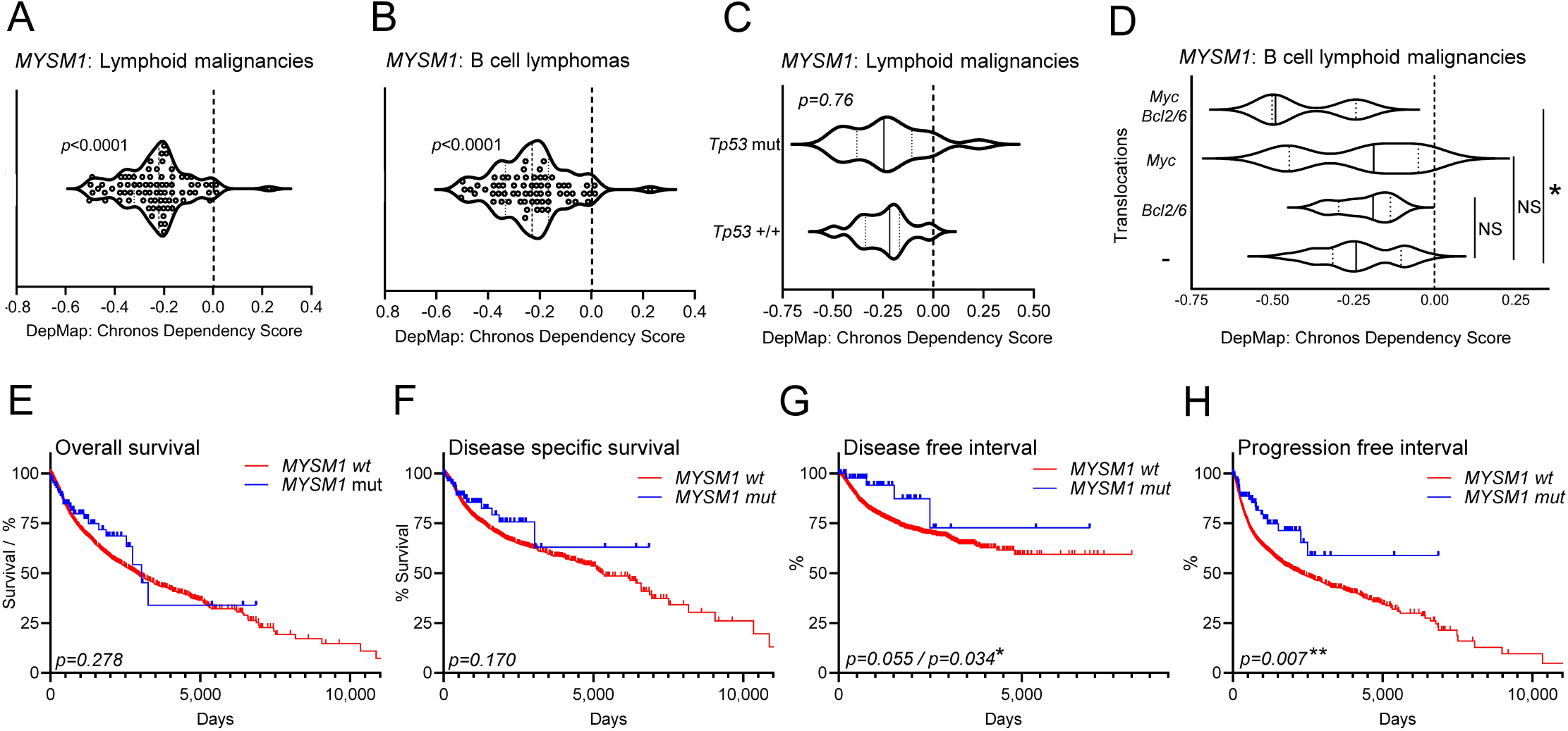
Protective effects of MYSM1 loss in human lymphoma. (A-D) Cancer Dependency Map Project database was interrogated for CRISPR-screen data for *MYSM1* gene in lymphoid malignancies (DepMap, https://depmap.org, 23Q4) (52). Chronos dependency scores were compared either between cell lines or to zero, which corresponds to the null hypothesis ‘no effect of MYSM1 loss on cell fitness’, using (A-B) one-sample *t*-test, (C) Student’s *t*-test, or (D) ANOVA with Dunnett’s post-hoc test. Significant impact of MYSM1 loss on cell fitness is demonstrated for **(A)** all human lymphoid cell lines in the database and **(B)** human B cell lymphoma cell lines, **(C)** independently of the *Tp53* status of the cells. **(D)** Double-hit or triple-hit lymphoma cell lines, characterized by *MYC*-locus translocations together with *BCL2* and/or *BCL6* translocations, are significantly more sensitive to *MYSM1* loss-of-function relative to other lymphoma cell lines. **(E-H)** The Cancer Genome Atlas (TCGA) was interrogated via cBioPortal (www.cbioportal.org) (53, 54, 55, 56) for correlation between *MYSM1* genotype and clinical prognosis. Kaplan–Meier survival curves comparing patient outcomes between tumors with non-silent somatic *MYSM1* mutations and *MYSM1* wild-type tumors, including **(E)** overall survival, **(F)** disease-specific survival, **(G)** disease-free interval, and **(H)** progression-free interval, for patients stratified by *MYSM1* mutation status across 32 non-redundant TCGA studies spanning a variety of cancer types. Tumors with non-silent *MYSM1* mutations (*n* = 91) are shown in blue, while *MYSM1* wild-type tumors (*n* = 8,943) are shown in red. Significance was assessed with log-rank Mantel-Cox and Gehan-Breslow-Wilcoxon tests; *p* values shown are from Mantel-Cox test, except for (G) where both *p* values are included due to the difference in outcome between the tests.

To analyze the correlation between *MYSM1* somatic mutations in human cancers and clinical outcomes, the Cancer Genome Atlas (TCGA) was interrogated via cBioPortal for relevant data across 32 non-redundant studies spanning many cancer types (cbioportal.org) (53, 54, 55, 56). This demonstrated that non-silent somatic *MYSM1* mutations in the tumors correlated with improved disease-free interval and progression-free interval for the patients, although without a significant effect on overall survival (Figure 5E-H). Furthermore, increased *MYSM1* copy number in the tumors correlated with inferior outcomes across all the metrics, including overall survival, disease-specific survival, disease-free interval, and progression-free interval (Figure S6C-F). Overall, this supports MYSM1 as a putative novel cancer drug target not only in mouse but also in human.

## DISCUSSION

In the current work, we established that a selective loss of MYSM1 DUB catalytic activity without the loss of MYSM1 protein expression is sufficient for protection against MYC-driven lymphoma in murine models, indicating MYSM1 as a putative drug-target. We characterized the effects of MYSM1 loss-of-function on both tumor cell physiology and antitumor immunity, demonstrating that MYSM1 loss within lymphoma cells strongly enhances immune cell infiltration into the tumors, while its selective loss in immune cells and the tumor microenvironment (TME) has complex modulating effects on immune cell infiltration, activation, and polarization, although no significant impact on disease progression or mouse survival.

Thus, the cell intrinsic loss of MYSM1 DUB catalytic activity in *EμMYC* cancer cells plays the major role in the protection against lymphoma progression, while its loss in the TME, though modulates immune cell infiltration, does not significantly impact the disease outcome. Mechanistically, our previous studies with non-cancerous *Mysm1*^Δ/Δ^ hematopoietic cells demonstrated reduced expression of the MYSM1 and MYC co-regulated genes encoding ribosomal proteins and translation factors, resulting in significantly decreased cellular protein synthesis rates (12). Analyses of the *EμMYC Mysm1*^Δ/DN^ lymphoma cells in our current study indicate that similar mechanisms also contribute to the protection against lymphoma with the loss of MYSM1 DUB catalytic activity. This suggests that the regulation of the MYC oncogenic transcriptional programs by MYSM1 requires its DUB catalytic function, and future studies will need to address whether such regulation operates primarily through the deubiquitination of histone H2AK119ub (11, 64, 65), which is a repressive epigenetic mark and a well-known substrate of MYSM1, or also through its activities on other alternative substrates.

Our study provides first evidence, from the DepMap and TCGA cancer genome databases, that the loss of MYSM1 impairs the fitness of human lymphoma cell lines in culture and correlates with some improvements in the clinical outcomes in cancer patients. Translational development of this work will require further validation of such protective effects and underlying mechanisms with human lymphoma models. Notably, MYSM1 proteins of human and mouse share ∼85% sequence homology, and there is a strong conservation of MYSM1 loss-of-function phenotypes between mouse models and rare human patients with congenital *MYSM1* mutations (OMIM: 618116) (11, 12, 66, 67). Furthermore, essential role of MYSM1 in the maintenance of MYC-regulated transcriptional programs shown in our previous work in mouse (12, 14) has been recently confirmed in human (13). We therefore anticipate that the loss of MYSM1 function in human lymphoma cells will show consistent effects with our studies in the murine *EμMYC* model and support MYSM1 as a novel drug-target for MYC-driven B cell lymphoma.

DUBs are promising drug-targets for many diseases, including cancer (15, 16), and the recent advances in the development of DUB inhibitors for cancer therapy are best exemplified by USP7 (68, 69, 70, 71). The current study demonstrates that an inducible loss of MYSM1 DUB catalytic activity in murine lymphoma provides strong protection against the disease (14), establishing a rationale for the development of inhibitors targeting MYSM1 as cancer therapeutics. MYSM1 belongs to the JAMM/MPN metalloprotease family of DUBs that rely on Zn^2+^ for catalytic activity (11, 72, 73). Drugs targeting other Zn^2+^ metalloproteases, such as matrix metalloproteases, have entered clinical trials (74). This suggests that the development of MYSM1 antagonists should be feasible, however inhibitor specificity may be a challenge due to similarity in catalytic mechanisms across the JAMM/MPN as well as other DUB family proteins (75). Structural data can be critical to identify the distinct modes through which DUBs bind to their substrates and inhibitors, and to exploit them for the optimization of inhibitor activities and specificities (76). MYSM1 comprises chromatin binding SANT and SWIRM domains, followed by the catalytic MPN domain (11), however only the structures of MYSM1 SANT and SWIRM domains have been published to date (77). We anticipate that high resolution structural data for MYSM1 MPN domain and full-length protein may facilitate the development of MYSM1 inhibitors as possible cancer therapeutics.

As a model of spontaneous cancer in an autologous and immunocompetent host, *EμMYC* mouse represents an ideal tool for mechanistic interrogation of both the tumor and immune functions in cancer therapy (78, 79, 80, 81). Oncogenic MYC is known to promote cancer immune evasion (19, 20, 21); and boosting antitumor immunity in *EμMYC* mice through tumor-based vaccination or checkpoint blockade can therefore prolong survival (82, 83, 84). Our study for the first time systematically interrogated the role of MYSM1 in antitumor immunity, revealing its distinct and competing effects in cancer cells, immune cells, and the TME. Thus, a selective loss of MYSM1 in lymphoma cells strongly enhanced immune cell infiltration into the tumors, which may contribute to its protective effects against the disease; and future studies should address the changes in tumor cell secretome that mediate these effects. In contrast, loss of MYSM1 in dendritic cells (DCs), myeloid leukocytes, T lymphocytes, or systemically in the TME did not alter disease progression, despite some notable modulation of immune cell infiltration and activation within the tumors. Thus, a selective loss of MYSM1 in DCs enhanced immune cell infiltration and Th1 T cell polarization within the tumors, while its systemic loss in the TME reduced immune infiltration, while also reducing the abundance of FOXP3^+^ Tregs and enhancing M1 macrophage polarization. Given the role of MYSM1 in non-malignant hematopoiesis (11, 30, 31, 85, 86), its systemic loss is expected to have some deleterious effects on immune cell production. Strategies for the targeting of MYSM1 inhibitors to the tumors may therefore be appealing to minimize their systemic effects on immune cell development and any downstream impact on antitumor immunity.

## CONCLUSION

In summary, our study establishes the chromatin binding DUB MYSM1 as a putative drug target for MYC driven lymphoma in murine models, provides the rationale for the future analyses of MYSM1 activities and mechanisms in human lymphoma models, and sets the stage for the development of inhibitors targeting its DUB catalytic activity.

## Ethics Approval

All experiments were in accordance with the guidelines of the Canadian Council on Animal Care and protocol MCGL-7643 approved by the McGill Animal Care Committee.

## Consent for Publication

Not applicable.

## Availability of Data and Materials

Data from this study are available within the paper and its Supplementary Materials. Requests for further information, resources, and reagents should be directed to the lead contact, Ana Nijnik. RNA-seq data in Figures 1 and S1 are available from the National Center for Biotechnology Information GEO database under accession number GSE310117, www.ncbi.nlm.nih.gov/geo/query/acc.cgi?acc=GSE310117. Data in Figures 5 and S6 were generated by analyzing the publicly available datasets from the Cancer Dependency Map Project (DepMap, https://depmap.org, 23Q4), DSMZ-German Collection of Microorganisms and Cell Cultures (www.dsmz.de), the American Type Culture Collection (ATCC, www.atcc.org), and the Cancer Genome Atlas (TCGA) – PanCancer Atlas, interrogated via cBioPortal (www.cbioportal.org).

## Authors’ Contributions

Experiments were performed by DS and VP, with the support and assistance of YL and NN and the supervision of AN. The study was designed and the manuscript written by AN, DS, and VP. FR and SH provided essential expertise, protocols, and training, as well as conducted a critical review of the results of the study.

## Acknowledgments

Flow cytometry was performed at the Cell Vision Core Facility of the McGill Life Sciences Complex, with the support of the Canadian Foundation for Innovation, and we thank Camille Stegen and Julien Leconte for training and assistance. We thank Catherine Gagné and other staff of the McGill Comparative Medicine and Animal Resources Centre (CMARC) for mouse colony management, and Rachel Kim, Orthy Aiyana, Na Young Cha, Andrew Yim, Gabriela Blaszczyk, and Connor Prosty for mouse genotyping. We acknowledge the contribution of Dr. Jerry Pelletier who recently passed away and has been instrumental during the initial development of this project.

---

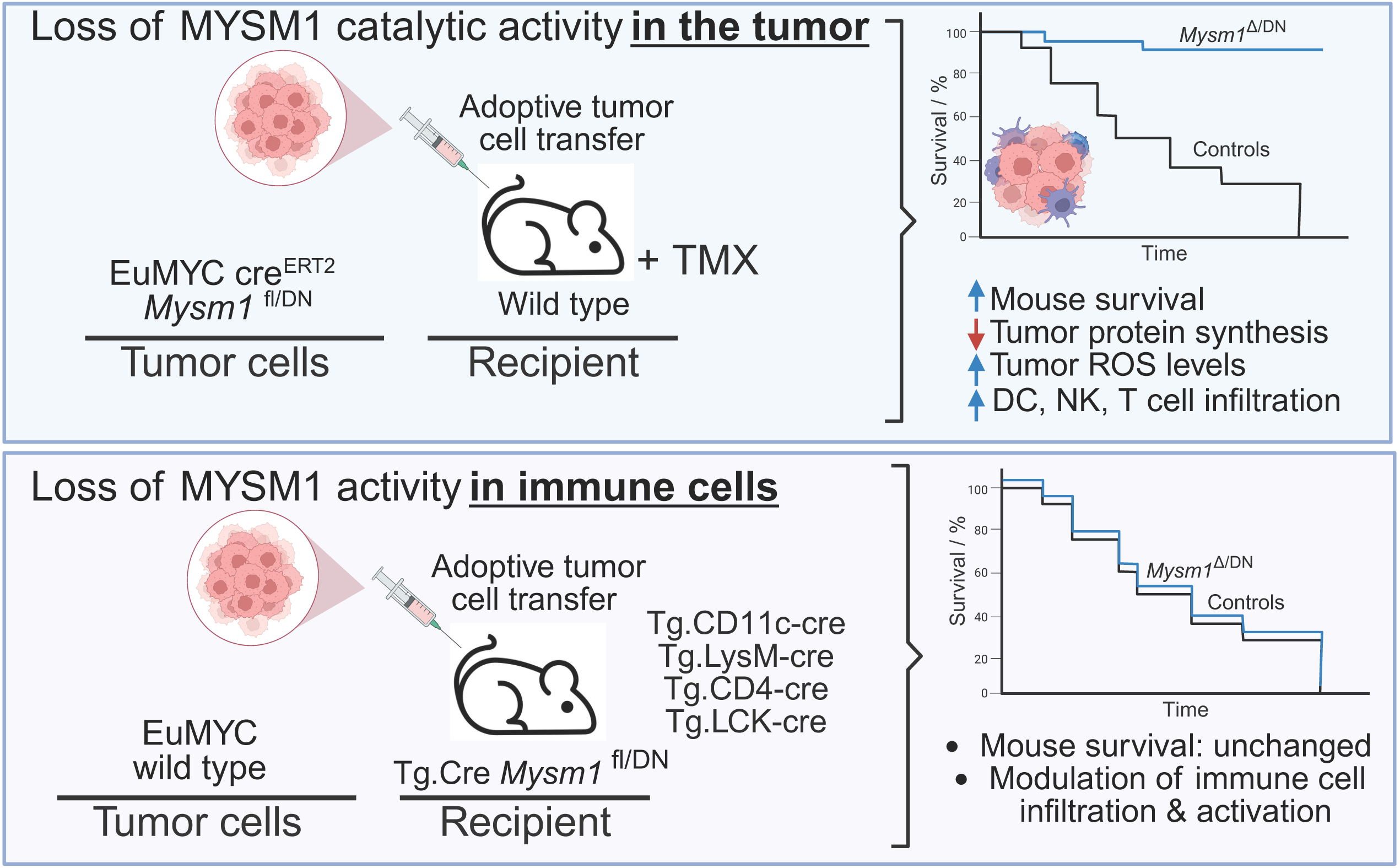

