## Supplemental Figures S1-S9, Supplemental Table S1, S5 for "Loss of MYSM1 deubiquitinase catalytic activity protects against MYC-driven B cell lymphoma via tumor intrinsic effects and indirect modulation of antitumor immunity"

**Figure S1. Loss of MYSM1 DUB catalytic activity: impact on the cancer cell transcriptome and physiology.** **(A-B)** Gene set enrichment analysis (GSEA) for enriched biological process terms in the bulk RNA-sequencing transcriptional data of *EμMYC* $Cre^{ERT2}$ *Mysm1*$^{\Delta/DN}$ versus control *EμMYC* $Cre^{ERT2}$ *Mysm1*$^{fl/DN}$ lymphoma cells, showing results for **(A)** pre-B lymphoma cells ($B220^{+}IgM^{-}$), and **(B)** mature B lymphoma cells ($B220^{+}IgM^{+}$). In both cases, *EμMYC* $Cre^{ERT2}$ *Mysm1*$^{\Delta/DN}$ (in vivo TMX-treated) lymphoma cells are compared to control *EμMYC* $Cre^{ERT2}$ *Mysm1*$^{fl/DN}$ (in vivo CO-treated) lymphoma cells, with negative normalized enrichment scores (NES) corresponding to downregulated transcriptional programs and positive NES corresponding to upregulated transcriptional programs. Select top biological process terms are labelled, with the full list provided in Supplemental Table S4. **(C)** *EμMYC* $Cre^{ERT2}$ *Mysm1*$^{fl/DN}$ tumor cells from TMX-induced or control corn oil treated mice were analyzed for immune activation and checkpoint markers by flow cytometry, gating on live $CD45^{+}B220^{+}$ lymphoma cells. Bars show mean ± SEM from n=10-11 tumor samples per genotype, consolidated from two independent experiments. Statistical analyses used Student's *t*-test; * $p$<0.05, *** $p$<0.001, and not significant if not indicated. **(D)** Representative flow cytometry histograms of CD86, CD80, CD40, MHCII and PD-L1, gated on live $CD45^{+}B220^{+}$ lymphoma cells, comparing across the *Mysm1* genotypes.

### Figure S1

#### Loss of MYSM1 catalytic activity in tumor cells

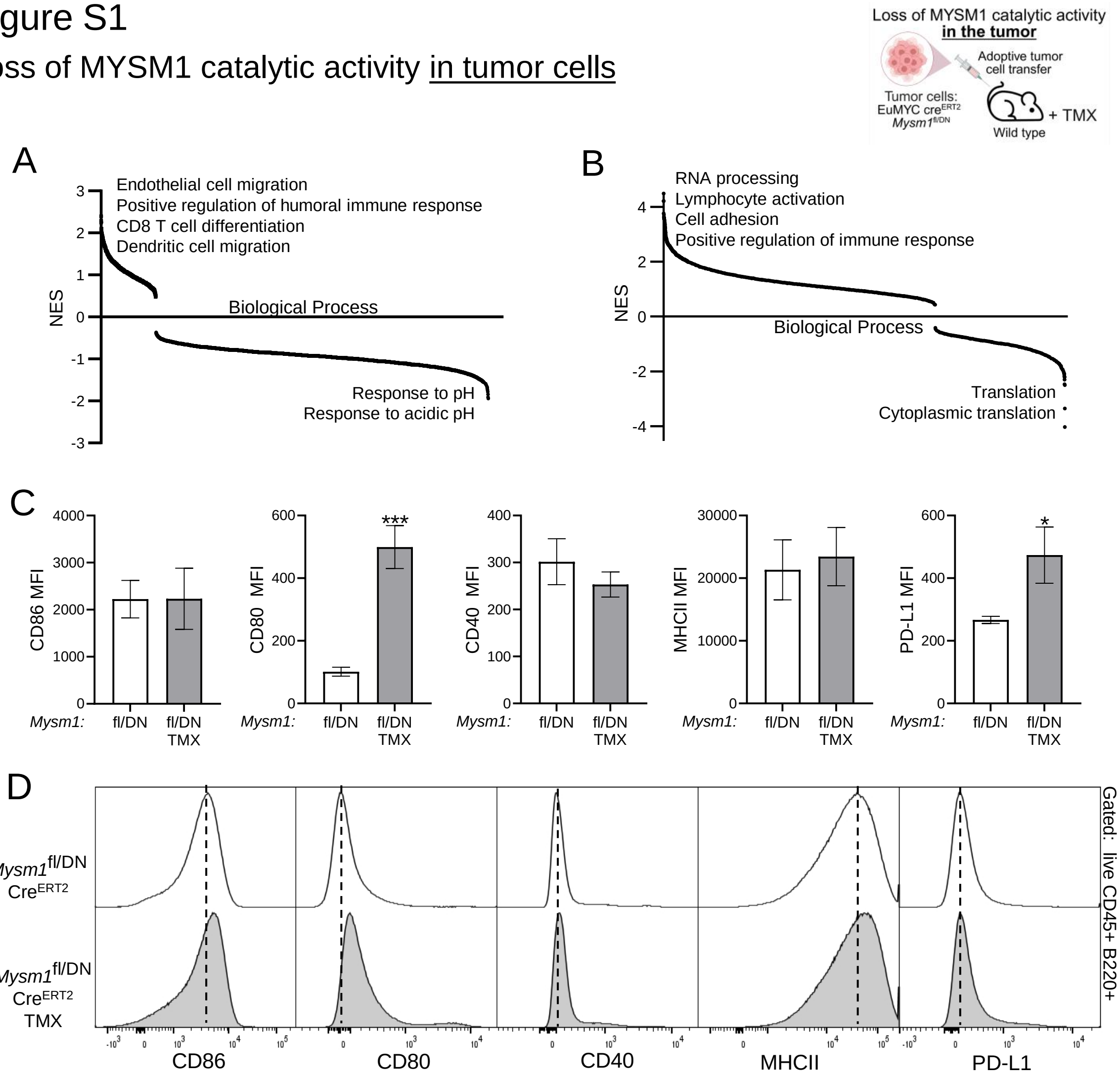

**Figure S2. Loss of MYSM1 catalytic activity in tumor cells indirectly modulates antitumor immunity.** *EμMYC* cells of *Mysm1*$^{fl/DN}$ Cre$^{ERT2}$ genotype were engrafted into wild type recipient mice and the mice treated with tamoxifen (TMX) to induce Cre$^{ERT2}$ activation or with corn oil as the vehicle control. Tumors were harvested from terminally ill mice for flow cytometry analyses. **(A-B)** Impact of the loss of MYSM1 catalytic activity in *EμMYC* lymphoma cells on the activation of tumor infiltrating dendritic cells: expression of CD86, CD80, CD40, MHCII, and PD-L1 evaluated by flow cytometry. Conventional dendritic cells (cDCs) were gated as live CD45$^{+}$B220$^{-}$Ly6G$^{-}$F4/80$^{-}$CD11c$^{+}$MHCII$^{+}$ cells, as shown in Figure S7. **(C-G)** Impact of the loss of MYSM1 catalytic activity in *EμMYC* lymphoma cells on tumor infiltrating T lymphocytes. Cells were gated as live CD45$^{+}$B220$^{-}$NK1.1$^{-}$CD3$^{+}$ followed by CD4$^{+}$CD8$^{-}$ or CD4$^{-}$CD8$^{+}$, as shown in Figure S8. **(C)** Expression of activation marker CD44 and exhaustion markers CTLA4 and PD1 on tumor infiltrating CD4 and CD8 T cells. **(D-E)** Proportion of CD4 T cells with positive intracellular staining for transcription factors Tbet, GATA3, RORγt, and FOXP3, corresponding to Th1, Th2, Th17 and Treg polarization, with isotype controls used to set the gates. **(F)** Representative flow cytometry plots showing the gating on CD4$^{+}$FOXP3$^{+}$ tumor infiltrating Treg cells. **(G)** Assessing the activation state of CD4 and CD8 T cells based on CD62L and CD44 marker expression, with CD62L$^{+}$CD44$^{-}$ corresponding to naive and CD62L$^{-}$CD44$^{+}$ to antigen experienced T cells. **(H-K)** Impact of the loss of MYSM1 catalytic activity in *EμMYC* lymphoma cells on the abundance and activation of myeloid cells within the tumors. Cells were gated as live CD45$^{+}$B220$^{-}$ , followed by Ly6G$^{+}$ for neutrophils, Ly6G$^{-}$Ly6C$^{+}$F4/80$^{-}$ for monocytes, Ly6G$^{-}$Ly6C$^{+}$F4/80$^{+}$ for monocyte derived macrophages, and Ly6G$^{-}$Ly6C$^{-}$F4/80$^{+}$ for tumor resident macrophages, as shown in Figure S9. **(H)** Monocytes, macrophages, and neutrophils quantified as a proportion of live cells within the tumor. **(I)** Representative flow cytometry plots showing increased abundance of monocytes and monocyte-derived macrophages in *Mysm1*$^{\Delta/DN}$ Cre$^{ERT2}$ tumors lacking MYSM1 catalytic activity. **(J-K)** Expression of activation markers CD86, CD80, CD40, MHCII and checkpoint marker PD-L1 on monocyte derived and tissue resident macrophages within the tumors. Numbers on plots indicate the percentage of cells within each gate out of the parent population, presented as mean ± SD for all the tumor samples of that experimental group. In all panels, bars represent mean ± SEM from n=10-11 tumor samples per group, consolidated from two independent experiments; MFI – mean fluorescence intensity. Statistical analyses used Student's *t*-test; * $p<0.05$ or not significant if not indicated.

### Figure S2

#### Loss of MYSM1 catalytic activity in tumor cells

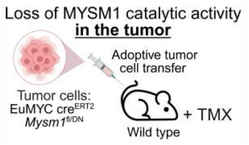

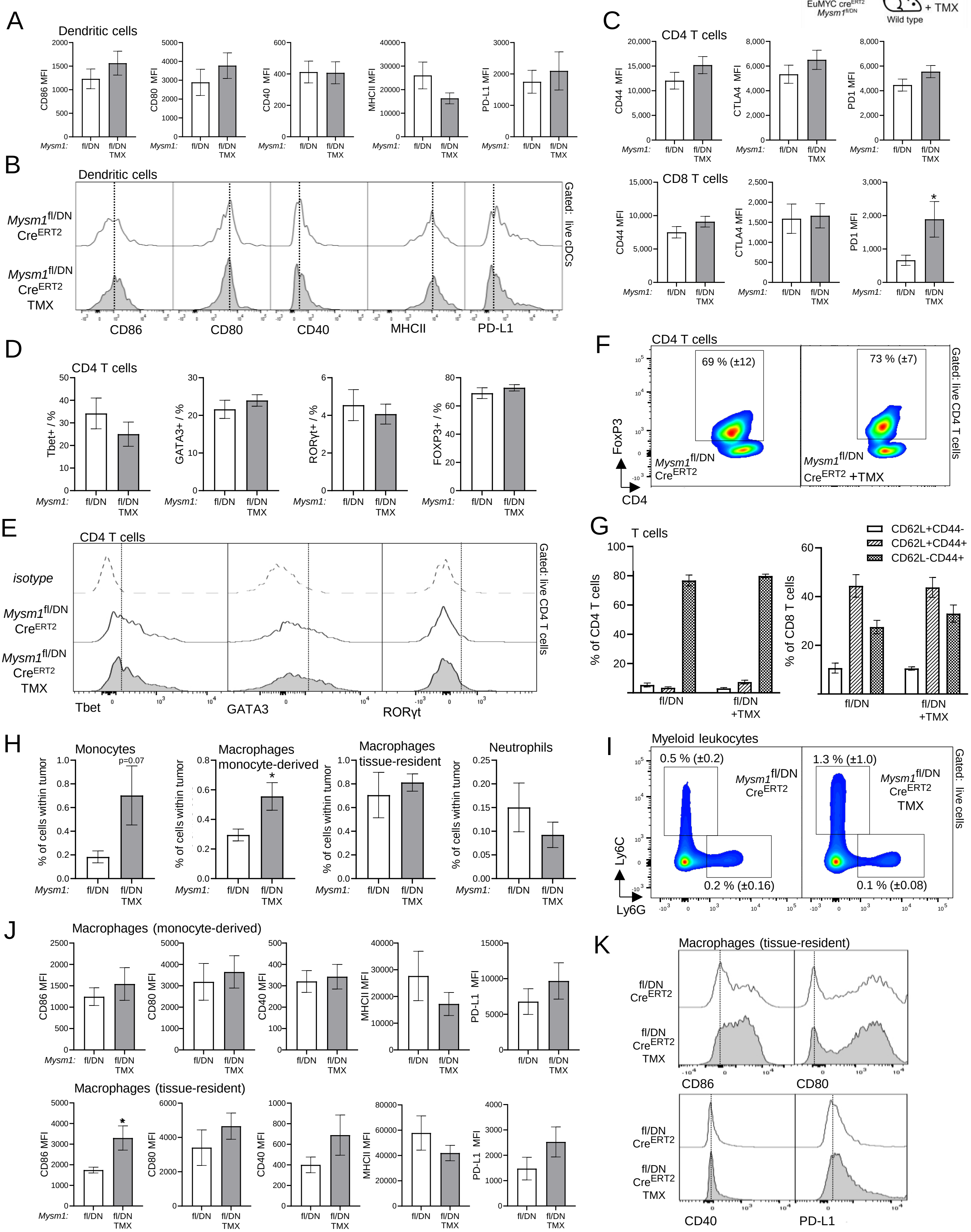

**Figure S3. Effects of the loss of MYSM1 function in dendritic cells or myeloid cells on immune cell infiltration and activation in the tumors.** Wild type *EμMYC* lymphoma cells were adoptively transferred into Cre transgenic recipient mice of *Mysm1*$^{DN/fl}$, *Mysm1*$^{fl/fl}$ and control *Mysm1*$^{fl/+}$, *Mysm1*$^{+/+}$ genotypes, tumors were harvested from terminally ill mice, and immune cell infiltration and activation was analyzed within the tumors. The following Cre-transgenic lines were used: CD11c-cre for *Mysm1* deletion in dendritic cells (DCs) (1, 2) and LysM-Cre for *Mysm1* deletion in myeloid lineage cells (3, 4). Dendritic cells (DCs) were gated as live $CD45^{+}Lin^{-}F4/80^{-}CD64^{-}$ $CD11c^{+}MHCII^{+}$ cells; myeloid cells were gated as live $CD45^{+}B220^{-}$ cells, followed by $Ly6G^{-}Ly6C^{+}F4/80^{+}$ for monocyte derived macrophages and $Ly6G^{-}Ly6C^{-}F4/80^{+}$ for tissue resident macrophages; CD4 T cells were gated as live $CD45^{+}B220^{-}NK1.1^{-}CD3^{+}CD4^{+}CD8^{-}$ cells. Bars represent mean ± SEM from n=3-10 tumor bearing mice per genotype (A-C) or n=5-8 tumor bearing mice per genotype (D); statistical analyses used Student's *t*-test; * $p<0.05$, *** $p<0.001$, and not significant if not indicated; MFI – mean fluorescence intensity. **(A-C)** Impact of the loss of MYSM1 function in dendritic cells on immune cell infiltration and activation within the tumors. **(A)** Activation state of tumor infiltrating DCs, as measured by CD86, CD80, CD40, MHCII and PD-L1 marker expression. **(B)** Proportions of tumor infiltrating CD4 T cells positive for transcription factor Tbet, corresponding to Th1 lineage, and representative flow cytometry histograms showing the staining of tumor infiltrating CD4 T cells for Tbet, with an isotype control antibody staining used to set the gates. **(C)** Activation state of tumor infiltrating monocyte derived macrophages, as measured by CD86, CD80, CD40, MHCII and PD-L1 marker expression. **(D)** Impact of the loss of MYSM1 function in myeloid cells on immune cell infiltration and activation within the tumors: activation state of tumor infiltrating monocyte derived macrophages and tissue resident macrophages is measured by CD86, CD80, CD40, MHCII and PD-L1 marker expression.

### Figure S3

#### Loss of MYSM1 activity in DCs and myeloid cells

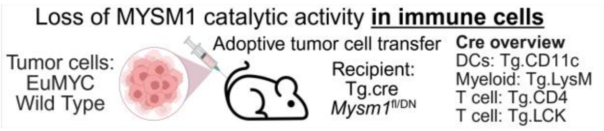

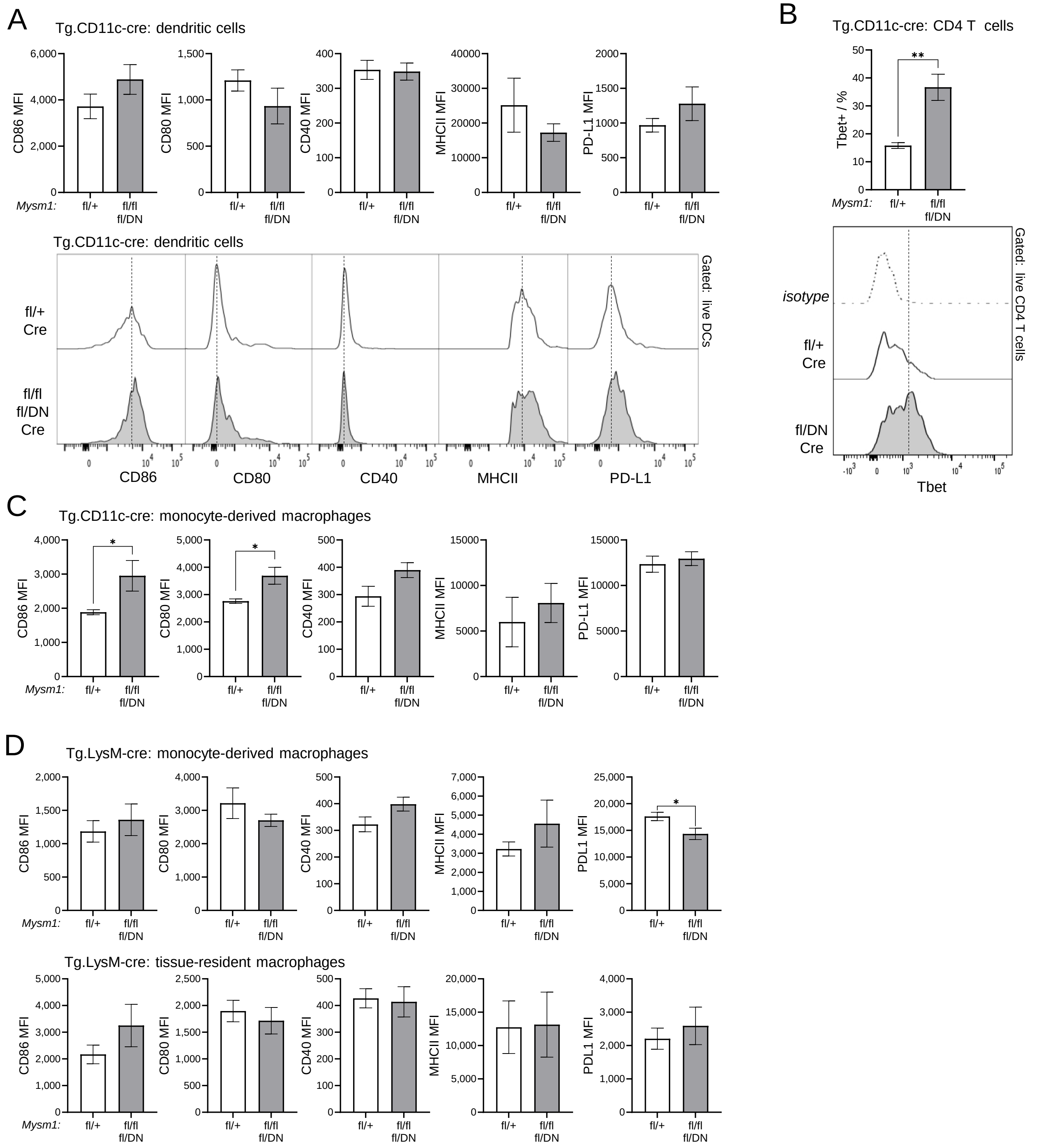

**Figure S4. Impact of the loss of MYSM1 in T lymphocytes on immune cell infiltration and activation in the tumors.** Wild type *EμMYC* lymphoma cells were adoptively transferred into CD4-cre transgenic recipient mice of *Mysm1*$^{fl/fl}$ and control *Mysm1*$^{+/+}$ genotypes (5, 6), tumors were harvested from terminally ill mice, and immune cell infiltration and activation was analyzed within the tumors. CD4 and CD8 T cells were gated as live CD45$^{+}$B220$^{-}$NK1.1$^{-}$CD3$^{+}$, followed by CD4$^{+}$CD8$^{-}$ or CD4$^{-}$CD8$^{+}$, respectively. Bars represent mean ± SEM from n=3-7 tumor bearing mice per genotype; statistical analyses used Student's *t*-test; results are not significant if not indicated. **(A)** Proportions of tumor infiltrating CD4 T cells with positive intracellular staining for transcription factors Tbet, GATA3, RORγt, and FOXP3 corresponding to Th1, Th2, Th17, and Treg lineages, respectively. **(B)** Representative flow cytometry histograms showing the staining of tumor infiltrating CD4 T cell for Tbet, GATA3, RORγt, and FOXP3, with isotype control antibody staining used to set the gates. **(C)** Activation state of CD4 and CD8 T cells based on CD62L and CD44 marker expression, with CD62L$^{+}$CD44$^{-}$ corresponding to naive T cells and CD62L$^{-}$CD44$^{+}$ to antigen experienced T cells.

### Figure S4

#### Loss of MYSM1 catalytic activity in T cells

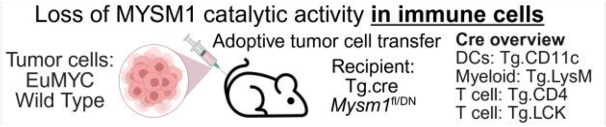

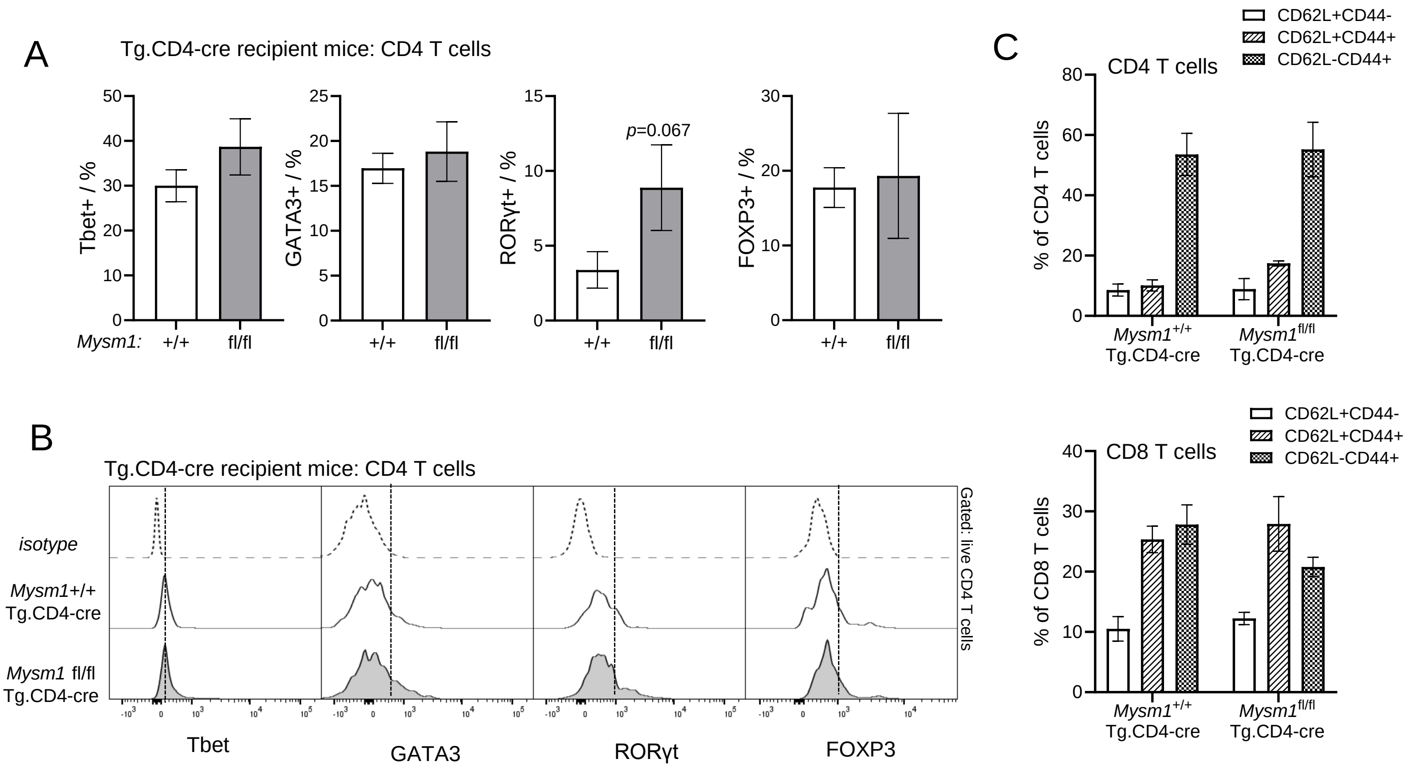

**Figure S5. Effects of a systemic loss of MYSM1 function in the microenvironment on the tumor infiltrating immune cells.** Wild type *EμMYC* lymphoma cells were adoptively transferred into $Cre^{ERT2}$ transgenic recipient mice for a systemic *Mysm1* deletion following tamoxifen (TMX) treatment. Tumor infiltrating immune cells are compared following TMX-treatment in cohorts of *Mysm1*$^{fl/fl}$ and *Mysm1*$^{fl/DN}$ mice against control *Mysm1*$^{fl/+}$ mice. Bars represent mean ± SEM from n=7-8 mice per group; MFI – mean fluorescence intensity. Statistical analyses used Student's *t*-test, * $p<0.05$, ** $p<0.01$, or not significant if not indicated. **(A-F)** Impact of a systemic loss of MYSM1 function in the tumor microenvironment on tumor infiltrating CD4 and CD8 T cells, gated as live $CD45^{+}B220^{-}NK1.1^{-}CD3^{+}$ cells followed by $CD4^{+}CD8^{-}$ or $CD4^{-}CD8^{+}$, respectively. **(A)** Proportions of tumor infiltrating CD4 T cells with positive intracellular staining for transcription factors Tbet, GATA3, and RORγt, corresponding to Th1, Th2, and Th17 lineages, respectively. **(B)** Representative flow cytometry histograms showing the staining of tumor infiltrating CD4 T cells for Tbet, GATA3, RORγt, with the isotype controls used to set the gates. **(C-D)** Expression of activation marker CD44 and exhaustion marker CTLA4 on tumor infiltrating CD4 and CD8 T cells. **(E-F)** Activation state of CD4 and CD8 T cells based on CD62L and CD44 marker expression, with $CD62L^{+}CD44^{-}$ corresponding to naive T cells and $CD62L^{-}CD44^{+}$ to antigen experienced T cells. **(G-J)** Impact of a systemic loss of MYSM1 function in the microenvironment on the activation of tumor infiltrating myeloid cells, gated as live $CD45^{+}B220^{-}Ly6G^{-}$ cells, followed by $Ly6C^{+}F4/80^{-}$ for monocytes, $F4/80^{+}$ for macrophages, and $F4/80^{-}CD11c^{+}MHCII^{+}$ for conventional dendritic cells (cDCs). Expression of activation markers CD86, CD80, CD40, MHCII and checkpoint marker PD-L1 on **(G-H)** monocytes, **(I)** macrophages, and **(J)** cDCs within the tumors.

### Figure S5

Systemic loss of MYSM1 catalytic activity in the tumor microenvironment

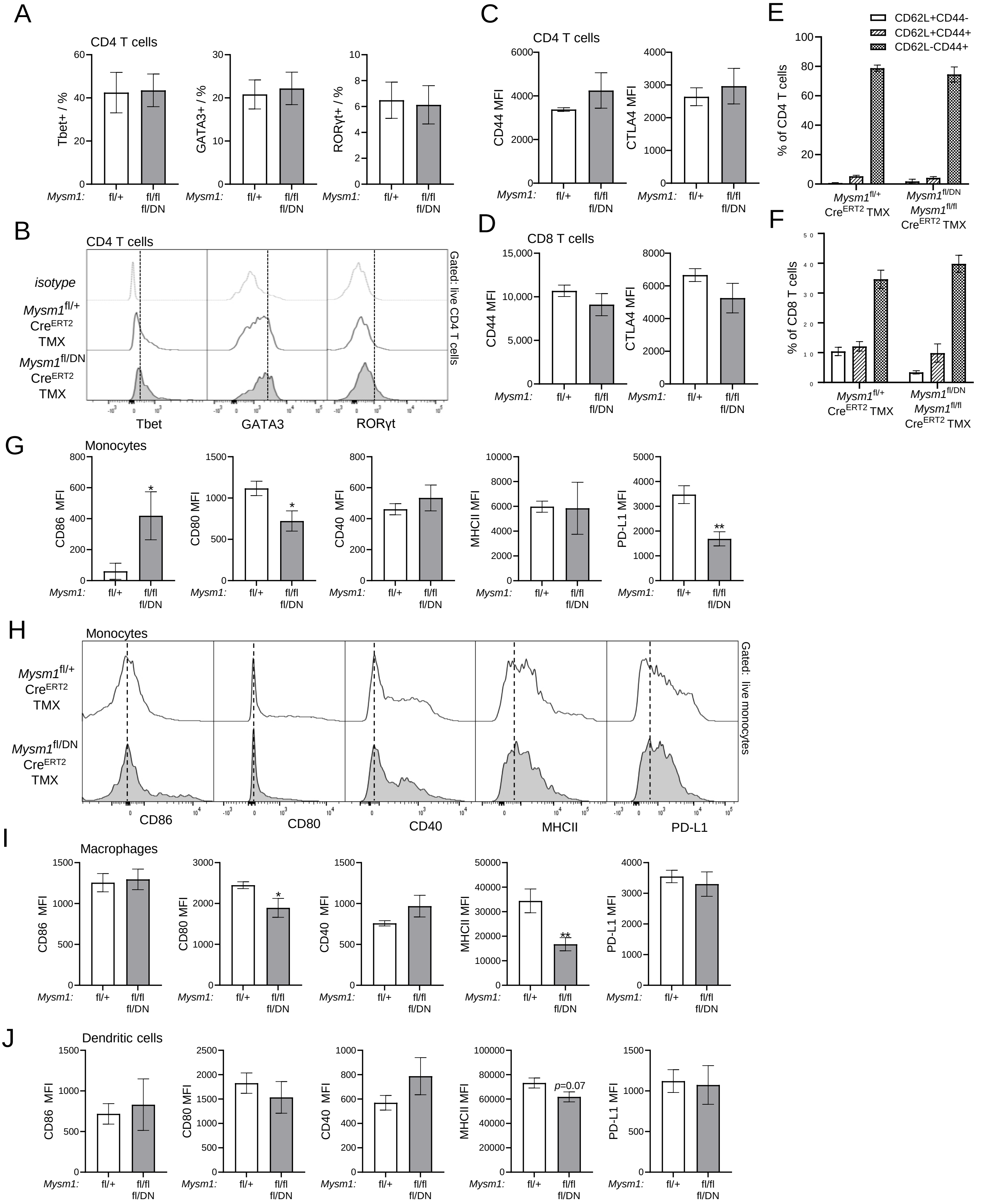

**Figure S6. Protective effects of MYSM1 loss in human lymphoma. (A-B)** Cancer Dependency Map Project database was interrogated for CRISPR-screen data for *MYSM1* gene and lymphoid malignancies (DepMap, *https://depmap.org*, 23Q4) (7). Information on the age and sex of the patients that the cell lines were derived from were obtained manually from the databases of DSMZ-German Collection of Microorganisms and Cell Cultures (*www.dsmz.de*) and the American Type Culture Collection (ATCC, *www.atcc.org*). Chronos dependency scores are compared between groups of human lymphoma cell lines based on **(A)** patient sex, and **(B)** patient age, analyzing with Student's *t*-test or test for linear trend in GraphPad Prism, respectively. **(C-F)** The Cancer Genome Atlas (TCGA) was interrogated via cBioPortal (*www.cbioportal.org*) (8, 9, 10, 11) for correlation between *MYSM1* genotype and clinical prognosis. Kaplan–Meier survival curves correlating *MYSM1* copy number with clinical outcomes, including **(C)** overall survival, **(D)** disease-specific survival, **(E)** disease-free interval, and **(F)** progression-free interval for patients stratified by *MYSM1* copy number across 32 non-redundant TCGA studies spanning a variety of cancer types. For copy number, gain represents a low-level gain of copy number whereas amplification indicates a higher-level gain of copy number, derived from copy-number analysis algorithms like GISTIC(12) or RAE(13). Statistical significance was assessed using the log-rank test (Mantel–Cox method).

Figure S6

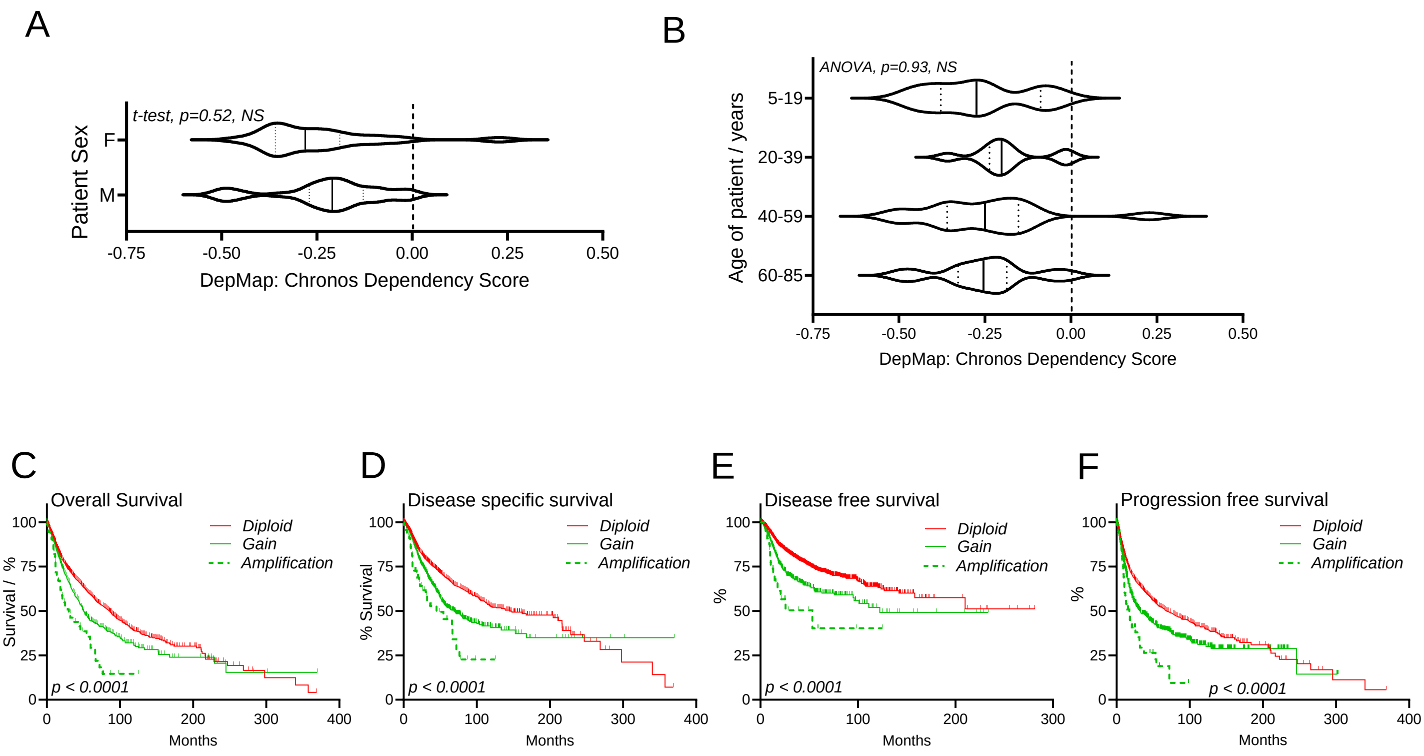
A
t-test, p=0.52, NS
Patient Sex
F
M
-0.75
-0.50
-0.25
0.00
0.25
0.50
DepMap: Chronos Dependency Score
B
ANOVA, p=0.93, NS
Age of patient / years
5-19
20-39
40-59
60-85
-0.75
-0.50
-0.25
0.00
0.25
0.50
DepMap: Chronos Dependency Score
C
Overall Survival
Diploid
Gain
Amplification
Survival / %
p < 0.0001
Months
D
Disease specific survival
Diploid
Gain
Amplification
% Survival
p < 0.0001
Months
E
Disease free survival
Diploid
Gain
Amplification
%
p < 0.0001
Months
F
Progression free survival
Diploid
Gain
Amplification
%
p < 0.0001
Months
0
25
50
75
100
0
100
200
300
400

**Figure S7. Flow cytometry gating scheme to characterize tumor infiltrating dendritic cells.** Lineage markers include CD3, CD19, NK1.1, Ly6G and TER119. DC – dendritic cells, cDC – conventional dendritic cells, pDC – plasmacytoid dendritic cells. All the antibodies used in the analyses are summarized in Table S1.

**Figure S8. Flow cytometry gating scheme to characterize tumor infiltrating NK cells and T lymphocytes.** T cells were subsequently analyzed for the expression of activation and exhaustion markers CD44, CD62L, CTLA4, LAG3, TIM3 and PD1, or for intracellular levels of transcription factors Tbet, GATA3, RORγt, and FOXP3. All the antibodies used in the analyses are summarized in Table S1.

**Figure S9. Flow cytometry gating scheme to characterize tumor resident and infiltrating myeloid cells, including macrophages, monocytes, and neutrophils.** The cells were subsequently analyzed for the expression of MHCII, activation markers CD86, CD80, CD40, and checkpoint marker PD-L1. All the antibodies used in the analyses are summarized in Table S1.

### Figure S7

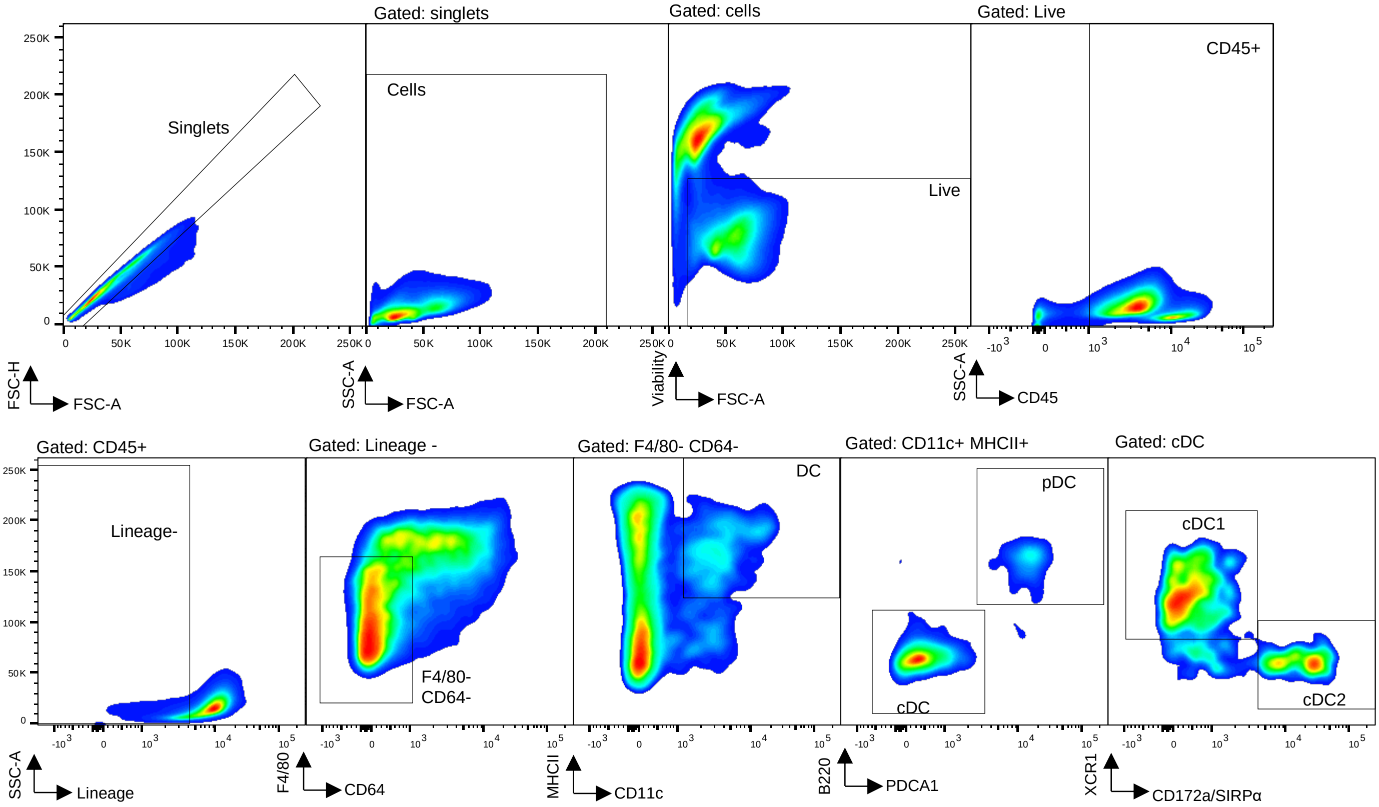

### Figure S8

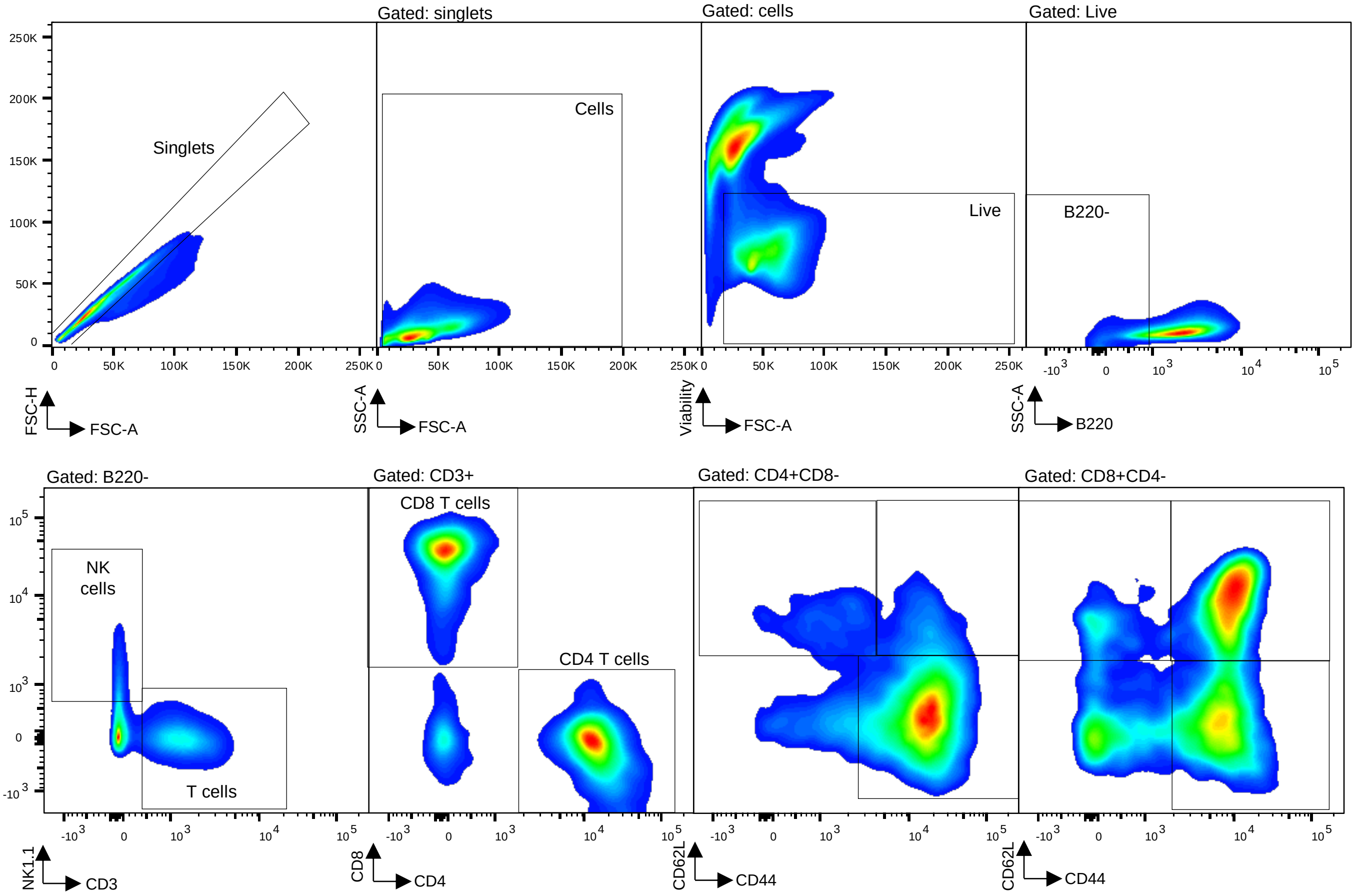

### Figure S9

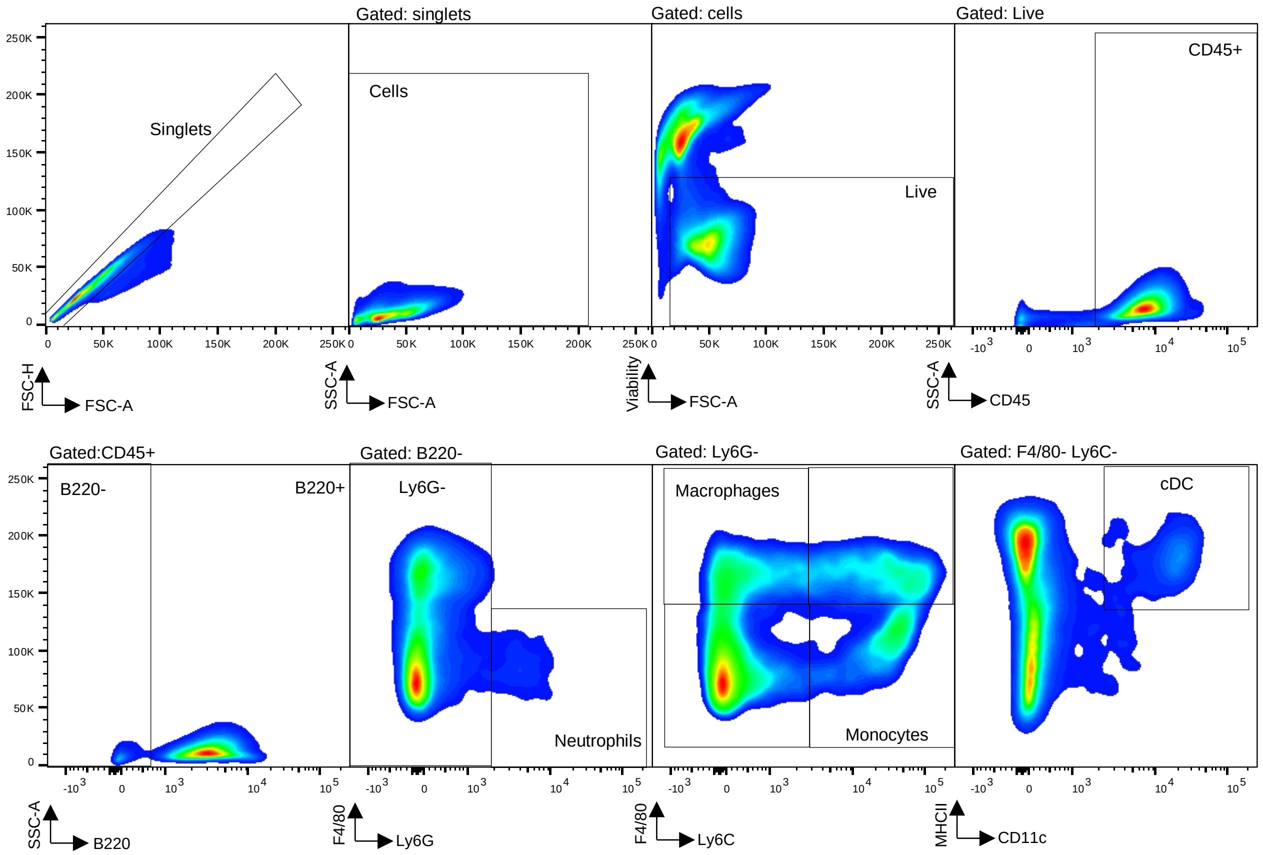
Gated: singlets
Gated: cells
Gated: Live
Singlets
Cells
Live
CD45+
FSC-H
FSC-A
SSC-A
FSC-A
Viability
FSC-A
SSC-A
CD45
Gated:CD45+
Gated: B220-
Gated: Ly6G-
Gated: F4/80- Ly6C-
B220-
B220+
Ly6G-
Neutrophils
Macrophages
Monocytes
cDC
SSC-A
B220
F4/80
Ly6G
F4/80
Ly6C
MHCII
CD11c

**Supplemental Table S1. Flow cytometry antibodies used in the study.**

| Product name | Clone | Supplier |
|---|---|---|
| **Panel 1 – Myeloid cells** | | |
| BUV395-conjugated anti-mouse CD45 | 30-F11 | BD Biosciences |
| eFluor450-conjugated anti-mouse CD45R/B220 | RA3-6B2 | Thermo Fisher Scientific |
| APC-Cy7-conjugated anti-mouse Ly6G | 1A8 | BioLegend |
| PerCPCy5.5-conjugated anti-mouse Ly6C | HK1.4 | BioLegend |
| BUV737-conjugated anti-mouse CD11c | N418 | BD Biosciences |
| BV650-conjugated anti-mouse MHCII | M5/114.15.2 | BioLegend |
| BV785-conjugated anti-mouse F4/80 | BM8 | BioLegend |
| PE-Cy7-conjugated anti-mouse CD274/PD-L1 | 10F.9G2 | BioLegend |
| PE-conjugated anti-mouse CD80 | 16-10A1 | Thermo Fisher Scientific |
| APC-conjugated anti-mouse CD86 | GL-1 | BioLegend |
| FITC-conjugated anti-mouse CD40 | HM40-3 | BioLegend |
| **Panel 2 – Dendritic cells** | | |
| BUV395-conjugated anti-mouse CD45 | 30-F11 | BD Biosciences |
| PerCPCy5.5-conjugated anti-mouse CD3 | 17A2 | BioLegend |
| PerCPCy5.5-conjugated anti-mouse CD19 | 6D5 | BioLegend |
| PerCPCy5.5-conjugated anti-mouse NK-1.1 | PK136 | BioLegend |
| PerCPCy5.5-conjugated anti-mouse Ly6G | 1A8 | BioLegend |
| PerCPCy5.5-conjugated anti-mouse TER119 | TER-119 | BioLegend |
| BUV737-conjugated anti-mouse CD11c | N418 | BD Biosciences |
| BV650-conjugated anti-mouse MHCII | M5/114.15.2 | BioLegend |
| BV785-conjugated anti-mouse F4/80 | BM8 | BioLegend |
| PE-Cy7-conjugated anti-mouse CD64 | X54-5/7.1 | BioLegend |
| eFluor450-conjugated anti-mouse CD45R/B220 | RA3-6B2 | Thermo Fisher Scientific |
| APC-Cy7-conjugated anti-mouse XCR1 | ZET | BioLegend |
| PE-conjugated anti-mouse CD172a/SIRPa | P84 | BioLegend |
| APC-conjugated anti-mouse CD371/CLEC12A | 5D3 | BioLegend |
| FITC-conjugated anti-mouse CD317/PDAC-1 | 927 | BioLegend |
| **Panel 3 – T cells and NK cells** | | |
| BUV395-conjugated anti-mouse CD3 | 145-2C11 | BD Biosciences |
| BUV737-conjugated anti-mouse CD4 | RM4-5 | BD Biosciences |
| BV650-conjugated anti-mouse CD8 | 53-6.7 | BioLegend |
| APC/eFluor 780-conjugated anti-mouse CD45R/B220 | RA3-6B2 | Thermo Fisher Scientific |
| PerCPCy5.5-conjugated anti-mouse NK-1.1 | PK136 | BioLegend |
| PE-conjugated, anti-mouse CD27 | LG.3A10 | BioLegend |
| eFluor 450-conjugated anti-mouse CD11b | M1/70 | Thermo Fisher Scientific |
| APC anti-human/mouse CD44 | IM7 | Thermo Fisher Scientific |
| PE-Cy7-conjugated anti-mouse CD279/PD1 | 29F.1A12 | BioLegend |
| BV785-conjugated anti-mouse CD223/LAG-3 | C9B7W | BioLegend |
| Alex-Fluor 488-conjugated anti-mouse CD366/TIM3 | 8B.2C12 | Thermo Fisher Scientific |

| | | |
|---|---|---|
| **Panel 4 – T cells** | | |
| BUV395-conjugated anti-mouse CD3 | 145-2C11 | BD Biosciences |
| BUV737-conjugated anti-mouse CD4 | RM4-5 | BD Biosciences |
| APC-Cy7-conjugated anti-mouse CD8a | 53-6.7 | BioLegend |
| PerCPCy5.5-conjugated anti-mouse/human CD44 | IM7 | BioLegend |
| BV650-conjugated anti-mouse CD45R/B220 | RA3-6B2 | BioLegend |
| BV785-conjugated anti-mouse CD62L | MEL-14 | BioLegend |
| PE-conjugated anti-T-bet | 4B10 | BioLegend |
| BV421-conjugated anti-mouse RORγt | Q31-378 | BD Biosciences |
| Alexa-Fluor 647-conjugated anti-GATA3 | 16E10A23 | BioLegend |
| FITC-conjugated monoclonal anti-FoxP3 | FJK-16s | Thermo Fisher Scientific |
| PE-Cy7-conjugated anti-mouse CD152/CTLA-4 | UC10-4B9 | BioLegend |
| **Panel 5 – Tumor cells and myeloid cells** | | |
| PerCPCy5.5-conjugated anti-mouse CD19 | 6D5 | BioLegend |
| eFluor450-conjugated anti-mouse CD45R/B220 | RA3-6B2 | Thermo Fisher Scientific |
| PE-Cy7-conjugated anti-mouse IgM | RMM-1 | BioLegend |
| BV785-conjugated anti-mouse F4/80 | BM8 | BioLegend |
| APC/Fire 750 conjugated anti-mouse CD64 (FcγRI) | X54-5/7.1 | BioLegend |
| APC conjugated anti-mouse CD163 | S15049I | BioLegend |
| FITC conjugated anti-mouse CD206 (MMR) | C068C2 | BioLegend |
| PE conjugated anti-Nos2 (iNOS) | W16030C | BioLegend |
| **Isotype Controls** | | |
| BV785-conjugated rat IgG1, $_K$ Isotype | RTK2071 | BioLegend |
| Alexa-Fluor® 488-conjugated rat IgG1, $_K$ Isotype | RTK2071 | BioLegend |
| PE-conjugated mouse IgG1 $_K$ Isotype | MOPC-21 | BioLegend |
| BV421-conjugated mouse IgG2a $_K$ Isotype | MOPC-173 | BioLegend |
| Alexa-Fluor® 647 mouse IgG2b $_K$ Isotype | MPC-11 | BioLegend |

**SUPPLEMENTAL TABLE LEGENDS (*all documents attached as excel files)***

**Supplemental Table S2. Bulk RNA-sequencing transcriptional analyses of *EμMYC* Cre$^{ERT2}$ *Mysm1*$^{Δ/DN}$ versus control *EμMYC* Cre$^{ERT2}$ *Mysm1*$^{fl/DN}$ lymphoma cells. (A)** Full list of genes expressed in pre-B B220$^{+}$IgM$^{–}$ lymphoma cells of *Mysm1*$^{Δ/DN}$ (tamoxifen treated, TMX) and control *Mysm1*$^{fl/DN}$ (corn oil treated, CO) genotypes. **(B)** List of genes differentially expressed in *EμMYC* Cre$^{ERT2}$ *Mysm1*$^{Δ/DN}$ (TMX) relative to control *Mysm1*$^{fl/DN}$ (CO) pre-B lymphoma cells. Significantly upregulated genes are at the top of the list, followed by significantly downregulated genes below. **(C)** Full list of genes expressed in B220$^{+}$IgM$^{+}$ mature B lymphoma cells of *Mysm1*$^{Δ/DN}$ (tamoxifen treated, TMX) and control *Mysm1*$^{fl/DN}$ (corn oil treated, CO) genotypes. **(D)** List of genes differentially expressed in *EμMYC* Cre$^{ERT2}$ *Mysm1*$^{Δ/DN}$ (TMX) relative to control *Mysm1*$^{fl/DN}$ (CO) mature lymphoma cells. Significantly upregulated genes are at the top of the list, followed by significantly downregulated genes below. **(B, D)** Differential gene expression analyzed at fold change (FC) ≥ |1.5| and false discovery rate (FDR) ≤ 0.001. Information provided for each gene includes gene name, fold change (FC), log2FC, *p*-value, false discovery rate (FDR), and counts per million (CPM). Ensembl gene ID and gene biotype are provided when available.

**Supplemental Table S3. Gene ontology (GO) analysis showing the enriched biological processes in the bulk RNA-sequencing transcriptional data of *EμMYC* Cre$^{ERT2}$ *Mysm1*$^{Δ/DN}$ versus control *EμMYC* Cre$^{ERT2}$ *Mysm1*$^{fl/DN}$ lymphoma cells. (A)** GO term analysis for significantly downregulated genes expressed in pre-B lymphoma cells (B220$^{+}$IgM$^{–}$). **(B)** GO term analysis for significantly upregulated genes expressed in pre-B lymphoma cells (B220$^{+}$IgM$^{–}$). **(C)** GO term analysis for significantly downregulated genes expressed in mature B lymphoma cells (B220$^{+}$IgM$^{+}$). **(D)** GO term analysis for significantly upregulated genes expressed in mature B lymphoma cells (B220$^{+}$IgM$^{+}$). In all cases, *EμMYC* Cre$^{ERT2}$ *Mysm1*$^{Δ/DN}$ (TMX-treated) cells are compared to control *EμMYC* Cre$^{ERT2}$ *Mysm1*$^{fl/DN}$ (CO-treated) cells, and differential gene expression is analyzed at fold change (FC) ≥ |1.5| and false discovery rate (FDR) ≤ 0.001.

**Supplemental Table S4. Gene set enrichment analysis (GSEA) for enriched biological process terms in the bulk RNA-sequencing transcriptional data of *EμMYC* Cre$^{ERT2}$ *Mysm1*$^{Δ/DN}$ versus control *EμMYC* Cre$^{ERT2}$ *Mysm1*$^{fl/DN}$ lymphoma cells. (A-B)** GSEA analysis of protein-coding genes expressed in pre-B lymphoma cells (B220$^{+}$IgM$^{–}$), with the terms with negative normalized enrichment scores (NES) corresponding to the downregulated transcriptional programs presented in **(A)**, and those with positive NES corresponding to the upregulated transcriptional programs presented in **(B)**. **(C-D)** GSEA analysis of protein-coding genes expressed in mature B lymphoma cells (B220$^{+}$IgM$^{+}$), with the terms with negative NES corresponding to the downregulated transcriptional programs presented in **(C)**, and those with positive NES corresponding to the upregulated transcriptional programs presented in **(D)**. In all cases, *EμMYC* Cre$^{ERT2}$ *Mysm1*$^{Δ/DN}$ (TMX-treated) lymphoma cells are compared to control *EμMYC* Cre$^{ERT2}$ *Mysm1*$^{fl/DN}$ (CO-treated) lymphoma cells.

**Supplemental Table S5. De novo promotor-based motif enrichment analysis for genes differentially expressed in *EμMYC* Cre$^{ERT2}$ *Mysm1*$^{Δ/DN}$ versus control *EμMYC* Cre$^{ERT2}$ *Mysm1*$^{fl/DN}$ lymphoma cells** (log2 FC >|1.5|, FDR <0.001), including **(A)** downregulated and **(B)** upregulated genes in pre-B B220$^+$IgM$^-$ lymphoma cells; **(C)** downregulated and **(D)** upregulated genes in mature B220$^+$IgM$^+$ lymphoma cells, analyzing within ±1,000 bp of the transcriptional start sites for each gene set.

| Motif | Name | *p*-value | %Peaks with motif (%background with motif) |
|---|---|---|---|
| **A) Motif enrichment analysis for downregulated genes in pre-B cell lymphoma** | | | |
| 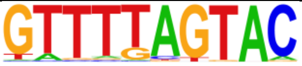 | HOXD12 | $10^{-11}$ | 8.4% (2.4%) |
| 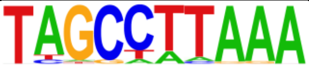 | CDX4/MA1473.2 | $10^{-10}$ | 10.5% (3.9%) |
| 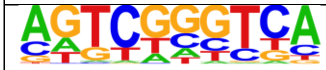 | RARA2 | $10^{-9}$ | 8.4% (2.4%) |
| 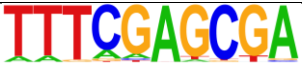 | E2F | $10^{-9}$ | 5.0% (1.2%) |
| 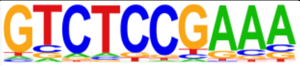 | MED 1 | $10^{-8}$ | 24.6% (14.7%) |
| **B) Motif enrichment analysis for upregulated genes in pre-B cell lymphoma** | | | |
| 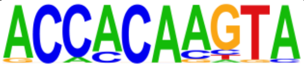 | RUNX2 | $10^{-10}$ | 21.9% (2.6%) |
| 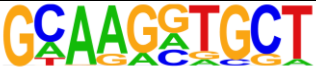 | IRF3 | $10^{-7}$ | 39.7% (13.4%) |
| 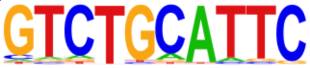 | POU5F1 | $10^{-7}$ | 28.8% (7.2%) |
| 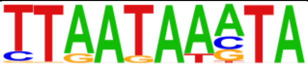 | HNF1b (Homeobox) | $10^{-7}$ | 19.2% (3.2%) |
| **C) Motif enrichment analysis for downregulated genes in mature B cell lymphoma** | | | |
| 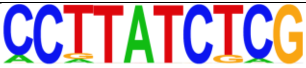 | GATA4 | $10^{-10}$ | 3.1% (0.4%) |
| 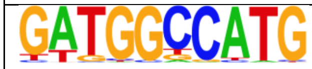 | SF1(NR) | $10^{-10}$ | 14.5% (6.8%) |
| 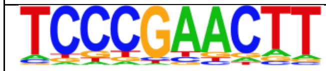 | NR1I3 | $10^{-9}$ | 26.8% (16.8%) |
| 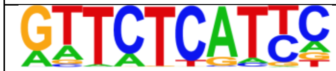 | POU6F1 | $10^{-8}$ | 7.9% (2.8%) |
| 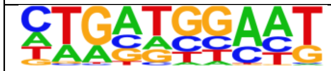 | ATF4 | $10^{-8}$ | 11.6% (5.2%) |
| **D) Motif enrichment analysis for upregulated genes in mature B cell lymphoma** | | | |
| 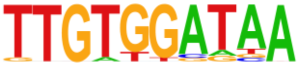 | FOXH1 | $10^{-8}$ | 27.54% (5.46%) |
| 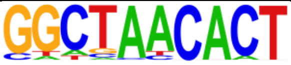 | SOX 1 | $10^{-8}$ | 15.94% (1.53%) |
| 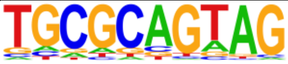 | OSR1 | $10^{-7}$ | 40.58% (13.85%) |
| 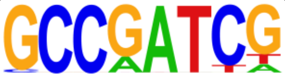 | NFY(CCAAT) | $10^{-7}$ | 31.88% (8.76%) |
| 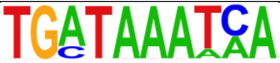 | PBX2 | $10^{-6}$ | 13.04% (1.29%) |
| 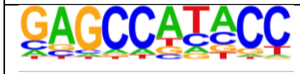 | KLF4 | $10^{-5}$ | 17.39% (3.12%) |
